# Spatial patterning regulates neuron numbers in the *Drosophila* visual system

**DOI:** 10.1101/2023.08.28.555170

**Authors:** Jennifer A. Malin, Yen-Chung Chen, Félix Simon, Evelyn Keefer, Claude Desplan

## Abstract

Neurons must be made in the correct proportions to communicate with the appropriate synaptic partners and form functional circuits. In the *Drosophila* visual system, multiple subtypes of Distal medulla (Dm) inhibitory interneurons are made in distinct, reproducible numbers—from 5 to 800 per optic lobe. These neurons are born from a crescent-shaped neuroepithelium called the Outer Proliferation Center (OPC), which can be subdivided into specific domains based on transcription factor expression. We fate mapped Dm neurons and found that more abundant neural types are born from larger neuroepithelial subdomains, while less abundant subtypes are born from smaller ones. Additionally, morphogenetic Dpp/BMP signaling provides a second layer of patterning that subdivides the neuroepithelium into smaller domains to provide more granular control of cell proportions. Apoptosis appears to play a minor role in regulating Dm neuron abundance. This work describes an underappreciated mechanism for the regulation of neuronal stoichiometry.

## Introduction

The function of any organ system depends on the existence of a roster of cell types present in reproducible proportions. Sensory systems such as the visual system illustrate this concept well: Adequately sampling the environment relies on information detected by photoreceptors, and interpretation of this information requires a specific number of interneurons at each intermediate layer of visual processing before it is sent to higher brain centers. Like the roughly 60 types of inhibitory amacrine cells of the mouse retina^1^, around 20 classes of *Drosophila* distal medulla (Dm) neurons exist and are made in distinct proportions (Figure 1A)^2–4^. The number of these neurons and the size of their arbors are essential for their function as they regulate the flow of visual information from the retina.

**Figure 1.**
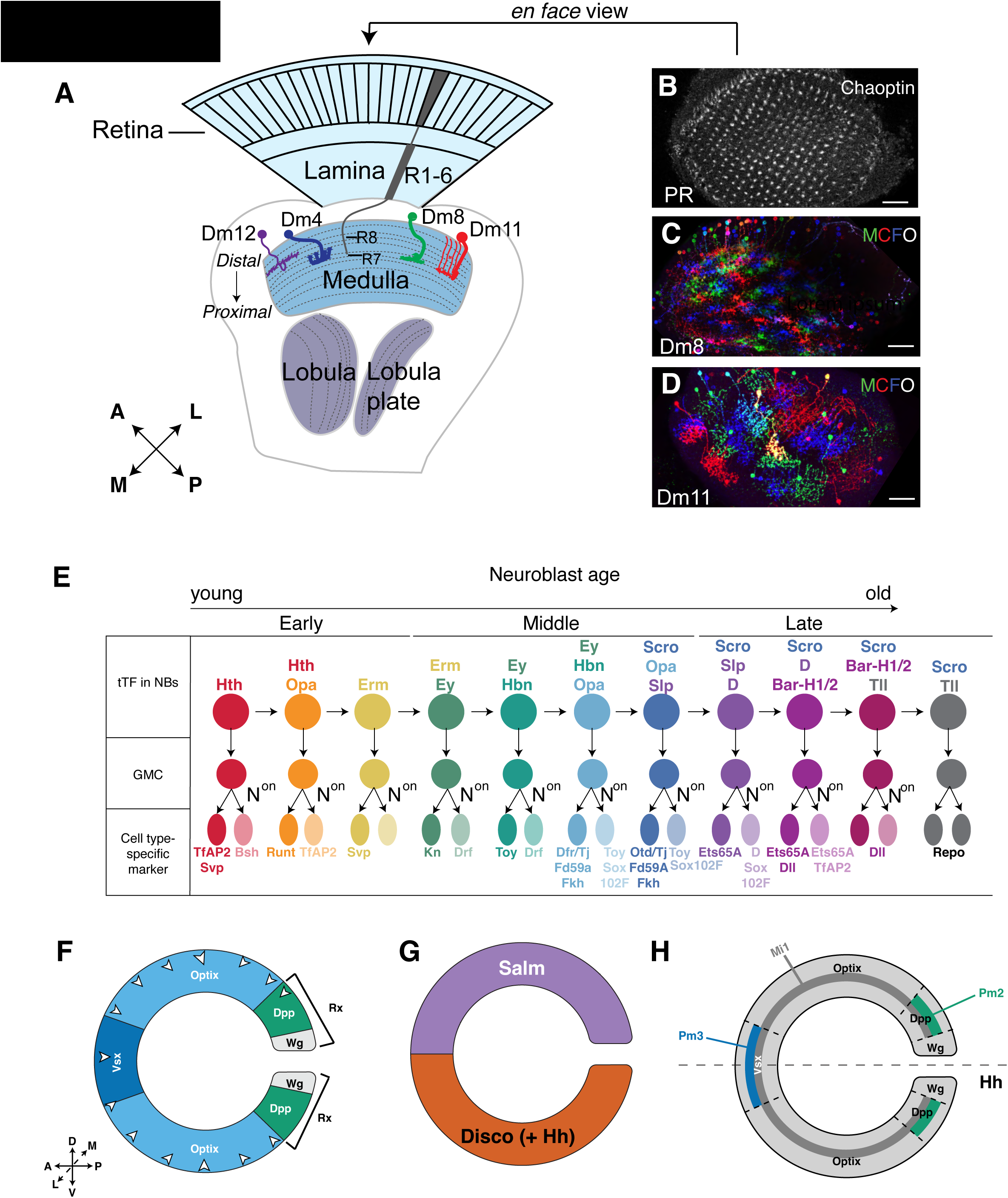
Anatomy and neuronal specification mechanisms of the optic lobe. (A) Graphical representation of the optic lobe with the retina and four neuropils: the lamina, medulla, lobula and lobula plate. The medulla contains over 100 cell types, roughly 20 of which are Distal medulla neurons. Some, like Dm8 (green), possess a 1:1 ratio of neurons to columns (800 cells per optic lobe); others, like Dm11 (red), are less numerous (70 cells per optic lobe). Dm4 has 40 neurons per optic lobe (blue), and Dm12 (purple) has 120. A= Anterior, P= Posterior, M= Medial, L= Lateral. (B) 800 photoreceptor ommatidia (stained by Chaoptin) project their axons into the medulla. Scale bar = 10μm. (C) Dm8 possesses roughly 800 cells per optic lobe. They are labeled by Multicolor Flip Out (MCFO), in which each neuron stochastically expresses a combination of HA, V5, and FLAG tags to individually label each cell type and their morphology. (D) Dm11 (also stained by MCFO) possesses roughly 70 cells per optic lobe. (E) Neural stem cells called neuroblasts express a series of temporal Transcription Factors (tTFs) as they age, the output of which directs neural patterning. N^on^= Notch^ON^, GMC= ganglion mother cell (Adapted from Konstantinides et al., 2022)^18^. (F) Medulla neurons are born from the Outer Proliferation Center (OPC), which is spatially subdivided based on the non-overlapping expression of transcription factors and growth factors^12^. D= Dorsal, V= Ventral, A= Anterior, P= Posterior, L= Lateral, M= Medial. Carats indicate the direction of neurogenic wave that produces neuroblasts. (G) The OPC can be dorsoventrally divided by the expression of Spalt and Disco^25^. (H) Fate mapping of medulla neurons born from the Hth temporal window^12^. Mi1s (gray), present at a 1:1 ratio of neurons to photoreceptor columns, are born from the entire main OPC. Pm2 (green) and Pm3 (blue) possess a small number of cells per optic lobe and are born from neuroblasts derived from smaller OPC subdomains.

Four neuropils comprise the *Drosophila* visual system: the lamina, the medulla, the lobula, and the lobula plate^5,6^. Dm neurons represent about one fifth of the 100 cell types in the medulla, the main optic ganglion through which visual information is processed *en route* to the central brain (Figure 1A). These Dm neurons are so called because they project their dendritic arbors to process visual information from the <u>d</u>istal half of the <u>m</u>edulla (Figure 1A)^2,5^. Dm neurons possess diverse morphologies and functions that are reflected in their distinct, yet stereotypic numbers. For example, highly overlapping Dm8 neurons receive input from R7 color-sensitive photoreceptors and are present at around 800 cells per optic lobe, *i.e.,* at a 1:1 ratio of neurons to photoreceptor unit-eyes, or columns (Figure 1A (green), Figure 1B-C)^2,7–9^. In contrast, Dm11 (Figure 1A (red), Figure 1D), the cell type that is most transcriptionally similar to Dm8, also receives inputs from R7s, but possesses only 70 tiled neurons per optic lobe^2,8,10^. How are these neurons, and others, generated with different proportions to achieve their distinct functions?

Dm neurons (and, in general, most medulla neurons) are born from a crescent-shaped neuroepithelium called the Outer Proliferation Center, or OPC^11,12^. During the third larval instar, a wave of differentiation passes through the neuroepithelium and, over a period of several days, converts the neuroepithelial cells into neural stem cells called neuroblasts^13–15^. Each neuroblast divides asymmetrically to regenerate a neuroblast and produce an intermediate progenitor (Ganglion Mother Cell, or GMC)^16^. During successive divisions, neuroblasts pass through a series of temporal windows generated by the overlapping expression of a cascade of temporal transcription factors (tTFs)—Hth, Opa, Erm, Ey, Hbn, Scro, Slp1/2, D, Bar-H1 and Tll—whose output specifies the fates of each different medulla neuron type (Figure 1E)^17–21^. Further cell fate diversification arises from the unique, asymmetric, Notch-mediated division of GMCs that generates two distinct daughter cells, one Notch^on^ and one Notch^off^ (Figure 1E)^17,22^. All OPC neuroblasts undergo the same temporal transcription factor series, with the exception of neuroblasts arising from the Wg spatial domain (*i.e.,* the tips of the OPC, or tOPC), which express a similar but modified series of tTFs^17,22^.

The output of the temporal series is modified by additional patterning generated by the expression of spatially restricted factors in the neuroepithelium. The OPC is partitioned into different domains based on the expression of various transcription factors and growth factors (Figure 1F-G)^12^. The visual system homeodomain transcription factor Vsx is expressed in the center of the OPC^23^; the Six3 homeodomain transcription factor Optix is expressed more posteriorly^24^, and the retinal homeodomain Rx is expressed at the tips of the OPC (Figure 1F)^12,22^. The OPC can be further subdivided along the dorsoventral axis: Spalt/Salm are expressed in the dorsal OPC while Disco is expressed ventrally (Figure 1G)^25^. The signaling molecule Hedgehog (Hh) is only expressed ventrally earlier in embryogenesis but is not maintained when mature neurons are formed (Figure 1G)^12,26^. The Rx domain is further subdivided into subdomains that are marked by the expression of Wingless (Wg) more posteriorly at the tips of the OPC, and Decapentaplegic (Dpp) closer to Optix (Figure 1F)^27,28^.

Previous experiments identified the neurons born from the earliest Hth temporal window: Mi1, a cell type with a 1:1 ratio of neurons to columns, is born from the entire OPC neuroepithelium, while less numerous cell types, such as Pm2 or Pm3, are born from much smaller neuroepithelial subdomains (Figure 1H)^12^. This suggested that spatiotemporal patterning could act not just as a mechanism for the regulation of cell fate, but that it could also regulate neuronal abundance. Although the spatial origins of neurons born from the Hth window are known, the spatial origins of later-born Dm neurons are unknown^12^. It is also unknown how much of a role spatial patterning plays for determining the proportion of different neurons.

We examined the role of spatial patterning in the regulation of Dm neuron cell number and cell fate specification using genetic fate mapping tools and single-cell RNA sequencing (scRNAseq) techniques. We find that the relative abundance of a specific Dm subclass is roughly proportional to the size of the neuroepithelium from which it is born. We identify additional spatial factors that pattern medulla neurons: Signaling from the BMP homolog Dpp provides additional spatial information via repression of the transcriptional repressor Brinker (Brk)^29^: Brk splits the Optix domain into subdomains. Furthermore, the overlap of Dpp expression with Optix forms an additional spatial domain within the neuroepithelium, allowing for more granular regulation of cell fate and cell number. We also show that apoptosis is not the main driver of cell number regulation and instead provides small, but significant changes to fine tune cell number.

Our work suggests that spatial patterning not only promotes cell fate, but that it also regulates cell type proportions.

## Results

### The number of Dm neurons is proportional to the size of their neuroepithelial domain of origin

As earlier experiments suggested that spatial patterning could regulate neuronal abundance^12^, we wondered whether the size of the neuroepithelial domain of origin correlated with relative abundance of Dm neurons. As spatial transcription factors are solely expressed in neuroepithelial cells and are not maintained in neuroblasts and mature neurons, we used genetic memory labeling techniques to map the spatial subdomain of birth for each Dm neuron.

To permanently mark neurons that are born from each spatial domain of interest, we crossed flies expressing an *actin::FRT-stop-FRT-*nuclear β−Gal cassette to flies expressing *UAS-Flp recombinase* under the control of a GAL4 line driven by the regulatory region of a spatially restricted factor (*e.g. Optix-GAL4*, Figure 2A)^9^. We then crossed our spatial factor fate mapping lines to lines in which GFP is specifically expressed in each neuron type of interest to determine its spatial origin in adult animals. As Dm neurons move during their development^12^, we could not use cell body position as a proxy for birth region during development. As the expression pattern of Rx is highly dynamic during development, we could not use it for our initial lineage tracing experiments (Supplementary Figure S1A-B). However, *Optix* and *hh* memory lines showed consistent expression in their respective domains throughout development, and so were used for further study^9^. These lines did not show additional expression later during development, and so did not require repression at other stages of development. We also used *pxb-GAL4* (a gene with the same OPC expression pattern as Vsx) to label the Vsx domain because Vsx is expressed outside of its domain in mature neurons, while Pxb is not^9,20^. The lineage tracing data and putative origins for each Dm line are presented in Figure 2C-N, as well as Supplementary Figure S1C-II, and is discussed in Table S1.

**Figure 2.**
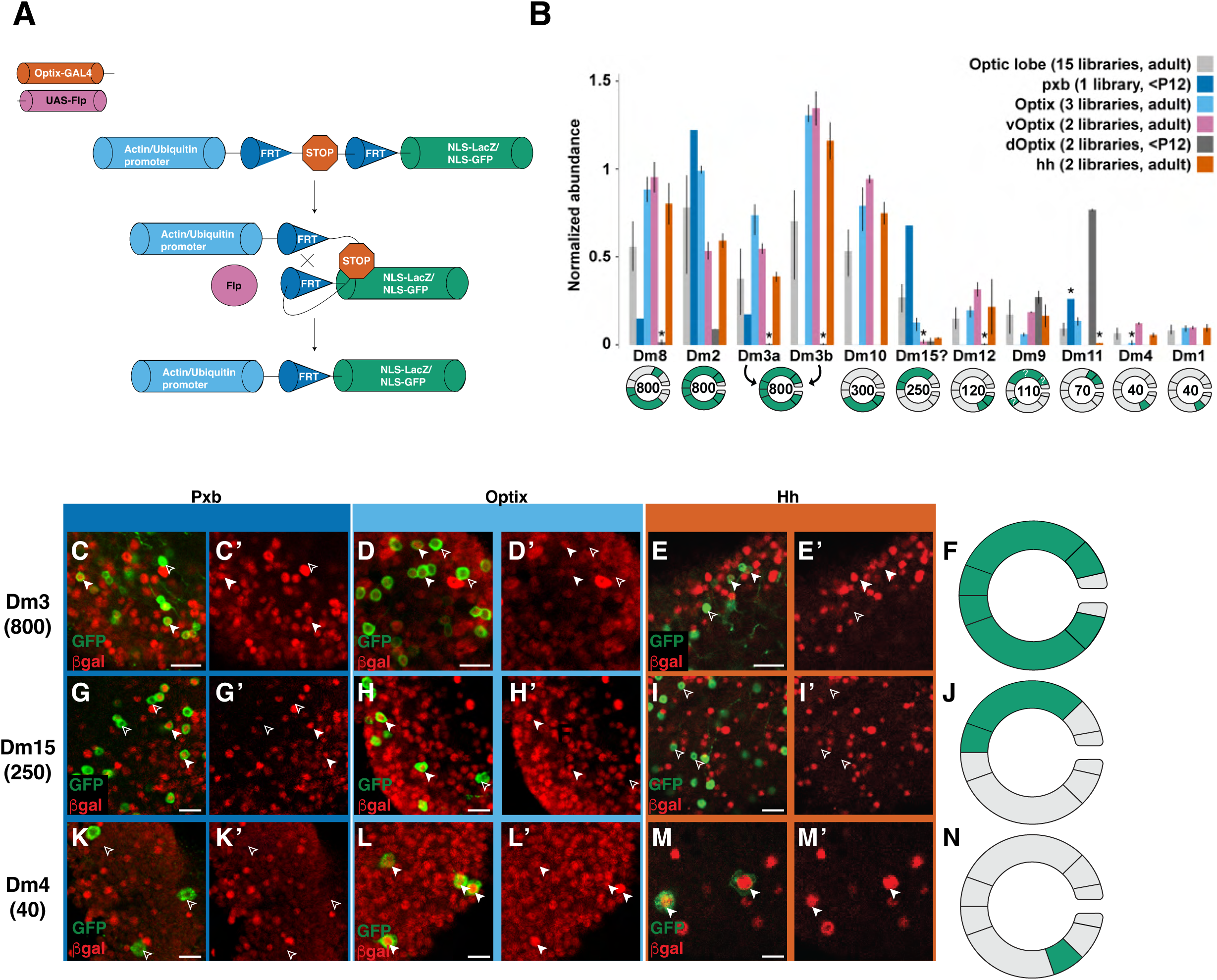
Fate mapping of Distal medulla neurons. (A) Technical approach for genetic fate mapping cassette^9^: A ubiquitious promoter (actin/ubiquitin) drives an FRT-Stop codon-FRT-nuclear localized reporter (lacZ or GFP); the stop codon is excised by expression of the Flp recombinase under the control of each spatial factor controlling GAL4. Neurons born from each neuroepithelial domain are thus permanently marked. (B) *pxb*, *Optix*, *vOptix*, *dOptix*, and *hh* lines were FACS sorted and subjected to scRNAseq to identify the spatial region of origin for each cell type. Average normalized abundance of the Dm clusters in datasets produced either in the whole optic lobe or from neurons from the FACSed datasets. The bars represent the minimal and maximal values across all libraries of a dataset. The asterisks indicate when at least 1 library had less than 3 cells of a given cell type, or if the annotations were made with low confidence. Below the graph represents the expected cell number and spatial origin of each cell type. This is based on the scRNAseq data as well as inferences made from all other experiments from the paper (see Table S1, Methods). (C-N) Representative example of spatial transcription factor fate mapping experiments. Scale bar = 5μm. Open carat= lack of expression. Filled carat = presence of expression. Dm3 neurons express the Pxb/Vsx lineage trace (C-C’), the Optix lineage trace (D-D’) and the Hh lineage trace (E-E’). (F) Graphical representation of Dm3 origin (based on information from Fig. 1B-E as well as Table S1). Dm15 neurons express the Pxb/Vsx lineage trace (G-G’), and the Optix lineage trace (H-H’), but none express the Hh lineage trace (I-I’). (J) Graphical representation of Dm15 origin (based on information from Fig. 1B, G-I as well as Table S1). Dm4 neurons do not express the Pxb/Vsx lineage trace (K-K’) but do express the Optix lineage trace (L-L’) and the Hh lineage trace (M-M’). (N) Graphical representation of Dm4 origin (based on information from Fig. 1B, K-M as well as Table S1).

Because the lineage tracing lines used GAL4 to express β-Gal, they required crosses to lines with cell-type specific enhancers from a different binary gene expression system (LexA rather than GAL4) to activate our GFP reporter line in specific Dm neurons. As a result, the lack of availability of specific LexA lines prevented us from fate mapping some of the Dm neurons. To obtain a more exhaustive description of Dm neuron origins, we used similar fate mapping lines crossed to *ubiquitin::FRT-stop-FRT-*nuclear GFP, which we sorted using FACS and submitted to single-cell RNA sequencing (scRNAseq). To distinguish between dorsal and ventral Optix, we used split-GAL4 lines that intersected Optix with the dorsal factor Spalt/Salm (*salr-T2A-VP16AD* **∩** *Optix-T2A-GAL4-DBD*) or the ventral factor Disco (*disco-T2A-VP16AD* **∩** *Optix-T2A-GAL4-DBD*, Figure 1G)^25^. For these experiments, we then used our existing scRNAseq datasets^18,30^ and a neural network classifier^30^ to accurately identify all clusters and their corresponding medulla neurons originating from each spatial domain (Figure 2B, Supplementary Figure S2A-C’; the results for neurons other than Dms will be presented in Simon et al., in preparation). As our dOptix line was crossed to a construct expressing GFP with a nuclear localization signal (*UAS-nls-GFP*) and not our lineage tracing line, we performed scRNAseq on animals who were sufficiently young that the GFP perdured in the maturing neurons (before P12). Some of these neurons were too immature to be accurately identified using the neural network classifier (*i.e.,* Dm2, Dm3, Dm15). However, when we constructed single-cell trajectories for immature neurons, we found that the precursors to Dm2, Dm3, and Dm15 were indeed detected in our FACSed dOptix datasets (Supplementary Figure S2A-C’, clusters Im11 and Im12).

The scRNAseq also identified the spatial origins of neurons belonging to unannotated clusters. To identify the Dm clusters among these data, we stained GAL4 lines expressing reporter genes for our neurons of interest with a panel of cell type-specific transcription factor markers^17,18^, which we then matched to clusters in our scRNAseq dataset (Supplementary Figure S3A-J’, Table S1). Although some cell types were too small in number to reliably annotate their clusters in the datasets, Dm15 was identified as a specific cluster (Supplementary Figure S4A-E).

Although the assignment to spatial domains were largely consistent between the individual fate mapping lines and the results of scRNAseq lines (Figure 2B, Supplementary Figure S1, S2), they also showed a few small differences between them (*i.e.,* some Dm11s came from Pxb (Vsx) in the scRNAseq, but we only identified a few of these neurons in our genetic lineage tracing experiment with *pxb-GAL4*, Figure 2B, Supplementary Figure S1C, Table S1). This could be due to contamination of the sequenced cell suspensions by unlabeled cells during FACSing, or mis-annotation of a single-cell transcriptome by our neural network. These caveats are discussed in Methods, Table S1 and in Simon et al. (in preparation).

We therefore looked at neurons where there was a discrepancy and immunostained them for transcription factor markers that we had identified for each Dm cell type in our scRNAseq datasets (Table S1, Supplementary Figures S5A-D). The larval location of these neurons within the medulla relative to the neuroepithelial domains allowed us to assign their origin more precisely. The reasoning used to assign the spatial origin of each Dm cell are explained in Table S1, as is the representation of the origin of each Dm subtype, which are also depicted in schematics in Figure 2B, F, J, N, Supplementary Figure S1 and Table S1.

Following these experiments, we found that the cell types represented in the highest number of lineage trace lines were the most numerous. For instance, Dm2 and Dm3 (∼800 neurons per optic lobe) originate from the entire dorsal and ventral main OPC (Figure 2B-F, Supplementary Figure S1C, Table S1). Dm8 was one exception as its 800 neurons come from a restricted ventral Optix domain (This exception is presented in the Discussion). Neurons in intermediate numbers (∼200-300) were found in fewer domains. For instance, Dm15s (∼250 neurons) only originate from the dorsal Vsx and Optix domains (Figure 2B, 2G-J, Supplementary Figure S1C, Table S1). Finally, neurons that were small in number, such as Dm4 (40 neurons per optic lobe) were born from single neuroepithelial subdomains (ventral Optix, Figure 2B, 2K-N, Supplementary Figure S1C, Table S1). However, the correlation between neuroepithelial domain size and neuron number became more tenuous with the less numerous neuron classes (Figure 2B). For example, there are roughly three times as many Dm12 (120 cells per optic lobe) neurons as there are Dm4s (40 cells) or Dm1s (40 cells); however, according to our immunofluorescence data (see “Additional OPC spatial subdivision is controlled by Brinker”), all appear to be born from the same subdomain within the ventral Optix domain (Figure 2B, 2K-N, Supplementary Figure S1C-G, S1X-AA, Table S1). We therefore wondered how cell number was regulated for less abundant cell types. We hypothesized that either different amounts of cell death determine the final differences in cell type numbers, or additional spatial patterning introduces finer compartments that differ in size, and the number of each neuron reflects size differences of these finer domains.

### Apoptosis plays a limited role in the regulation of Dm cell number

Programmed cell death has long been implicated in the regulation of neuron number across various nervous systems^31^. In the vertebrate nervous system, neurons are generated in excess, and those that do not receive sufficient neurotrophic support are removed via programmed cell death. In fact, as many as half of the neurons born in the vertebrate central nervous system are culled through programmed cell death^32^. In the *Drosophila* visual system, some Dm neurons are also made in excess and pared back during development^9,33,34^. To determine the role of apoptosis in the regulation of Dm cell number, we inhibited programmed cell death using a p35 baculovirus repeat domain protein transgene^35^ expressed under the control of a pan-neuronal enhancer (*elav-GAL4*) and determined whether the number of Dm neurons in adult animals increased. Seven of the 9 Dms tested did not show a significant difference in cell number following overexpression of p35, while the change in number was significant but modest for two neurons, Dm4 and Dm11 (Figure 3A, see Methods for quantification method). For example, Dm11s showed an increase from an average of 52 to 78 neurons after p35 overexpression (Figure 3A, P= 0.01, T-Test). Similarly, Dm4s showed a significant increase from 35 to 45 (Figure 3A, P=0.005, T-Test). To confirm that apoptosis was indeed not the main regulator of Dm cell number, we used other methods to inhibit cell death and measured the number of Dm4s: we performed RNAi against the effector caspase *dronc*^36^, expressed a dominant negative form of *dronc*^37^, and expressed microRNAs targeted to the upstream cell death activators *hid, reaper* and *grim*^38^. These mutations led to similar results to p35 overexpression (Figure 3B). The small changes observed corroborate the observation that some Dm/medulla neuron types are born in excess and undergo programmed cell death preceding synaptogenesis^9,33,34^, but are too modest to explain the differences between neurons with the same spatial origin (*e.g.,* Dm4 vs Dm12). Therefore, we suggest that apoptosis is mostly used for fine tuning of cell number, and that the initial setpoint of cell number regulation occurs earlier.

**Figure 3.**
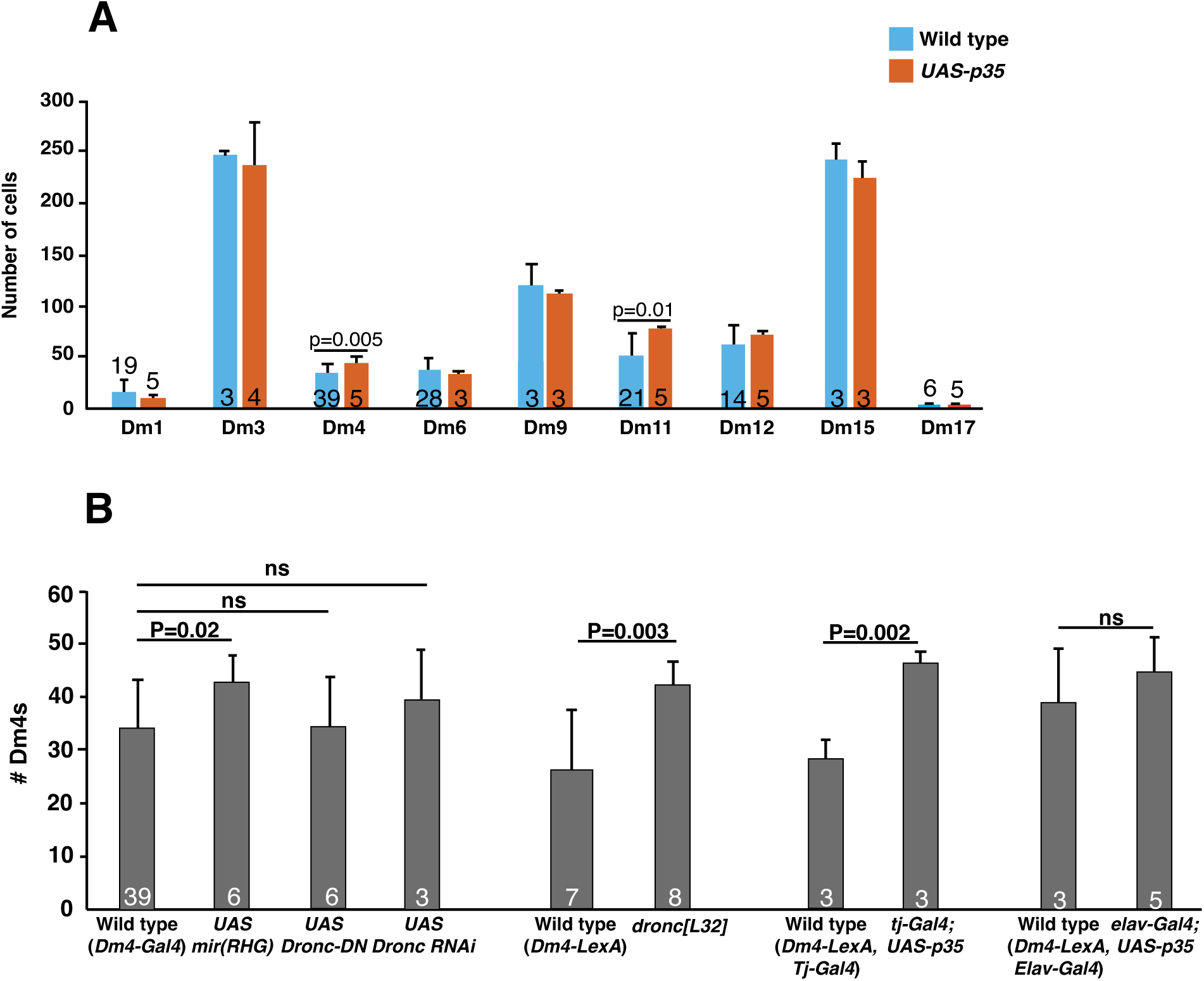
Apoptosis plays a minor role in regulating Dm neuron number. (A) Inhibition of apoptosis via expression of *elav-GAL4*; *UAS-p35* transgenes provides limited increases in cell number. Number within bars= number of animals scored, Error bars: standard deviation, T-test. (B) Inhibition of apoptosis via miRNA against *rpr*/*hid*/*grim*, *dronc RNAi*, *dronc-DN*, or *dronc*[L32] mutants leads to limited increases in Dm4 number. Error bars: standard deviation, small number within bars= number of animals scored, T-test. ns = not significant.

### Additional OPC spatial subdivision is controlled by Brinker

Previous experiments suggested that the OPC is further subdivided into smaller spatial subdomains^9^. For example, Dm8 neurons labelled with the transcription factors Dachshund (Dac) and Traffic Jam (Tj) were mapped to be born in the anterior 2/3 of the ventral Optix domain^9^. Furthermore, Dm8s exist as two distinct subtypes that differ in their connectivity to different classes of R7 color photoreceptors; pale Dm8s (pDm8) connect to pale R7s and are born from the anterior 1/3 of the ventral Optix region near the Vsx domain, while yellow Dm8s (yDm8) connect to yellow R7s and are born from the central 1/3 of the ventral Optix domain (Figure 4A-B’, Supplementary Figure S6A-A’)^9^. We therefore sought to identify the neurons born from the posterior 1/3 of the ventral Optix domain (closer to Dpp) and the mechanisms determining these subdomains. Dm1, Dm4 and Dm12, which express the transcription factors SoxN and Tj, were found in the posterior 1/3 of the ventral Optix domain (Figure 4A-B’, Supplementary Figure S6B-B’’). Thus, together, pDm8, yDm8 and Dm1/Dm4/Dm12 are born in at least three distinct spatial domains that cover the entirety of the ventral Optix domain (Figure 4A-B’).

**Figure 4.**
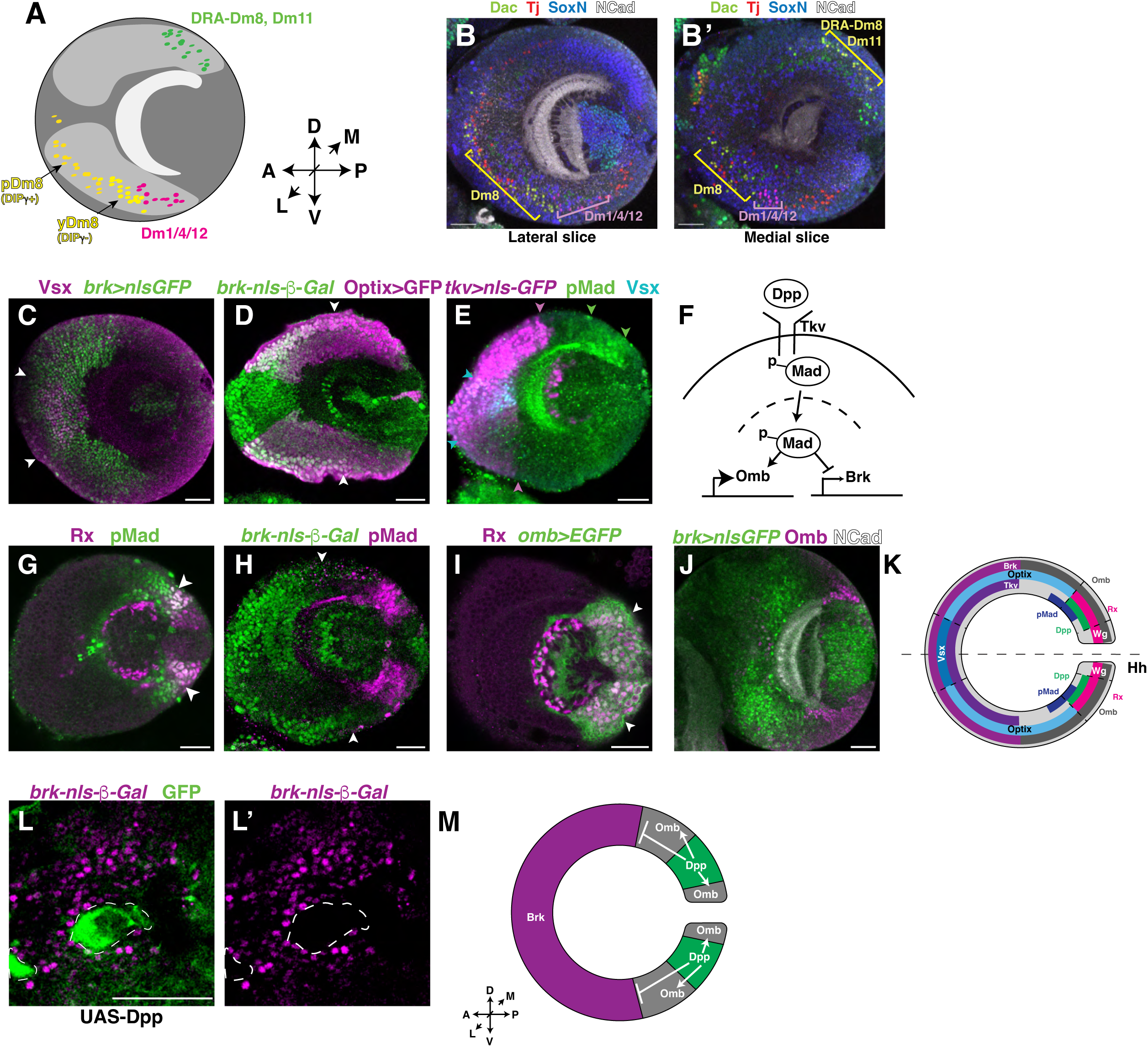
Mutually exclusive *brinker* and *dpp* expression delineates a second spatial patterning axis. (A) Model for spatial subpatterning of yDm8, pDm8, DRA-Dm8, Dm11, Dm1, Dm4 and Dm12 within the Optix domain (See also Table S1. Dm8 data from Courgeon and Desplan, 2019^9^). Lightest gray domain: medulla neuropil; Medium gray domain: Optix region; Darkest gray domain: rest of optic lobe. D= Dorsal, V= Ventral, A= Anterior, P= Posterior, L= Lateral, M= Medial. (B-B’) Dac+Tj+ y/pDm8s are born from the anterior 2/3 of the ventral Optix domain (a few Dac+ Tj+ cell bodies are also found in the medial Vsx domain), while SoxN+Tj+ Dm1/4/12 neurons are born from the posterior 1/3 of the ventral Optix domain. (B) Lateral slice, (B’) Medial slice. (C) Vsx expression sits completely within the *brk-GAL4*; *UAS-nls-sfGFP* OPC domain. Carat: border of Vsx OPC domain. (D) *brk*-nulacZ is expressed within the anterior 2/3 of the Optix region. Carat: border of Brk expression. (E) The Type I Receptor *tkv* (*tkv-GAL4*; *UAS-nls-GFP*) is expressed in a domain that is similar to *brk* expression (see Figure 4C). Magenta carat: border of Tkv expression, green carat: border of pMad expression, cyan carat: border of Vsx OPC expression. (F) Graphical representation of the Dpp signaling pathway. (G-J) Immunostaining experiments suggest that Dpp signaling pathway components are expressed in the third larval instar OPC in a manner similar to previously described systems. (G) pMad is expressed at the edge of the Rx domain, and into the Optix domain. Carat: region of overlap. (H) *brk-*nuLacZ and pMad share mutually exclusive expression patterns in the OPC. Carat: edge of Brk expression. (I) Omb is expressed within the Rx domain, and slightly outside. Carats: region of Omb/Rx overlap. (J) Omb expression sits adjacent to Brk. (K) Graphical representation of Dpp component expression patterns. (L-L’) *brk-nulacZ* expression is disrupted in *UAS-dpp* overexpression Flp-out clones. Dotted lines: region of disrupted *brk-nulacZ* expression where Dpp is overexpressed. (M) Model for relationship between Dpp and Brk signaling. Scale bar: 30μm.

The finding that multiple neuron types with different numbers span the ventral Optix domain suggested that additional spatial factors are used to pattern the medulla^9^. We found that two members of the BMP/Dpp pathway, the BMP/Dpp type I receptor Thickveins (Tkv) and the transcriptional repressor Brinker (Brk), were expressed in a domain spanning the entire Vsx domain and the anterior 2/3 of the Optix domain where Dm8s are produced (Figure 4C-E)^39^. Brk was therefore an excellent candidate to define the boundary between Dm8 and Dm1/Dm4/Dm12 in the ventral Optix domain. Co-staining these cells with Brk showed that Dm8 neurons end at the border of Brk expression (Supplementary Figure S6C-C’). In contrast, most of the Dm1s, Dm4s and Dm12s sat on the other side of the Brk boundary (Supplementary Figure S6D-D’). This suggests that Brk separates Dm8 from Dm1, Dm4 and Dm12 in the ventral OPC.

Similarly, two other classes of neurons, DRA-Dm8s and Dm11s, are born from the posterior 1/3 of the dorsal Optix domain closer to the Dpp domain (Figure 4A-B’, Supplementary Figure S6E-E’)^9,10,40^. However, we could not conclusively identify the neurons born from the anterior 2/3 of the dorsal Optix domain that overlaps with Brk.

### Dpp signaling establishes the Brinker-expressing domain in the OPC

As Brk forms an additional spatial subdomain within the OPC, we wondered how its expression pattern was established. Dpp is a BMP family protein that forms a morphogen gradient in the *Drosophila* wing and leg imaginal discs^41^. Dpp binding to its Type I receptor (Thickveins) leads to a signal transduction cascade ending with the phosphorylation of the transcription factor Mad (pMad). pMad activates multiple targets such as the gene encoding the transcription factor *optomotor blind* (*omb,* also known as *bifid*; Figure 4F)^42^ but also inhibits the expression of numerous genes, including *brk*, whose expression, in turn, represses genes dependent on Dpp^29^. We therefore tested whether the Brk domain was established by Dpp signaling at a distance from its expression domain. Indeed, pMad was expressed in the Rx (and thus, Dpp) domain and extended into the Optix domain, consistent with its activation by Dpp (Figure 4G). As with the wing/leg discs, *brk* was expressed adjacent to pMad (Figure 4H). Tkv was also expressed in a pattern mutually exclusive to pMad, similar to Brk expression (Figure 4E). In the leg disc, this allows for increased Dpp signaling where its receptor concentration is lowest^43^. As in the wing/leg discs^42^, Omb was expressed in response to active Dpp signaling: across the entire Rx domain, which includes both Dpp and Wg (Figure 4I), and as well as into the Optix domain where it bordered Brk expression (Figure 4J). Thus, the expression patterns of Dpp and its downstream targets suggest that Dpp acts in the OPC to activate pMad to define the domain of Brk expression (Figure 4K).

To assess whether Dpp represses Brk expression to delimit the size of its domain, we generated sparse Flp-out clones using *tubulin-*GAL4. Clones overexpressing Dpp and the surrounding region where Dpp diffuses lacked *brk-lacZ* expression, suggesting that Dpp suppresses *brk* expression to establish additional spatial domains within the OPC (Figure 4L-M).

### Brk and Dpp signaling alters the numbers of Dm neurons

Although Dpp and Brk expression are correlated with the positions of different neuron classes, are Dpp and Brk necessary and sufficient to regulate the abundance of the neurons born in their domain of activity? As Dm8 and Dm1/4/12 cells abut each other during larval development (Figure 4B-B’), we hypothesized that the gradient of Dpp that regulates Brk expression delineates the border between the cell types generated in these domains. Indeed, *dpp* overexpression under the control of an *Optix-GAL4* driver reduced the number of Dac^+^Tj^+^ Dm8 cells but increased the numbers of SoxN^+^Tj^+^ Dm1s, Dm4s, and Dm12s, indicating that it is required for the specification of Dm1/Dm4/Dm12 at the expense of Dm8 (Figure 5A-C’, Dm1/4/12: Dm8 ratio changed from 0.58 to 2.72, N= 3, P= 0.0001, Chi-square test). The remaining Dm8 cells were restricted to a domain closer to the Vsx domain (Figure 5C, yellow bracket). In contrast, *Optix-GAL4; UAS-dpp RNAi* increased the number of Dm8 cells and reduced the number of Dm1, Dm4 and Dm12 cells (Figure 5A, D, D’, Dm1/4/12: Dm8 ratio changed from 0.58 to 0.24, N= 3, P= 0.0001, Chi-square test). *Optix-GAL4; UAS-brk* larvae similarly exhibited a decrease in the number of Dm1/4/12 neurons and an increase in Dm8s (Figure 5A, E, E’. Dm1/4/12: Dm8 ratio changed from 0.58 to 0.24, N= 3, P= 0.0001, Chi-square test), while *Optix-GAL4*; *UAS-brk RNAi* showed a decrease in the number of Dm8 neurons with an increase in the proportion of Dm1/4/12s (Figure 5A, F, F, Dm1/4/12: Dm8 ratio changed from 0.58 to 2.15, N= 6, P= 0.0001, Chi-square test). Thus, Dpp and Brk act in an opposite manner to specify Dm8 vs. Dm1/4/12 fate and stoichiometry in subdomains of Optix.

**Figure 5:**
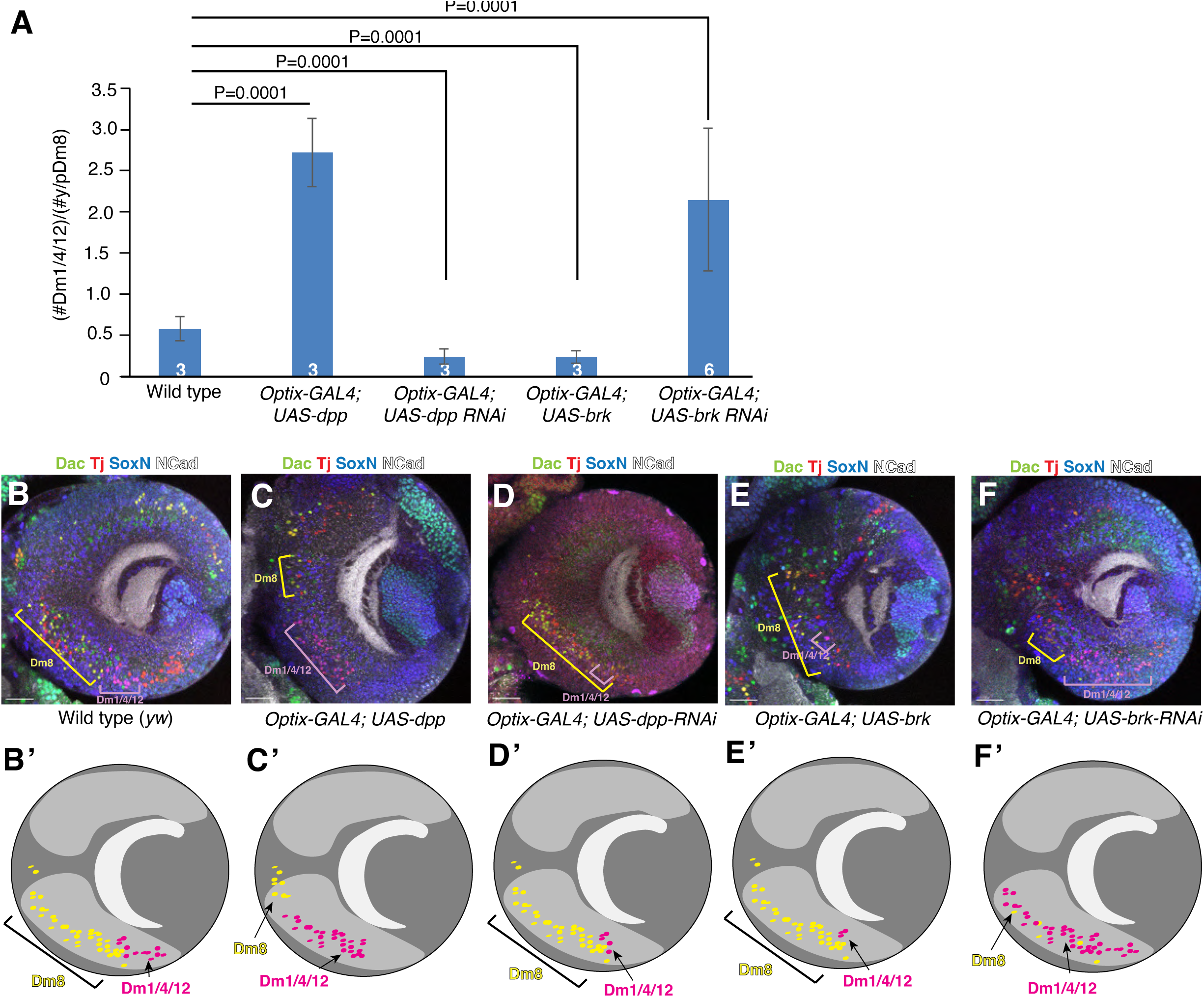
*brk* or *dpp* overexpression during larval development impacts Dm neuron number. (A) Quantification of images. Error bars = standard error of the mean. Small number inside bars= number of optic lobes scored. P value calculated using Chi Square test. (B) Wild-type Dac+Tj+ Dm8 and SoxN+Tj+ Dm1/4/12 neurons. (B’) Graphical representation of (B); Lightest gray domain: medulla neuropil; Medium gray domain: Optix region; Darkest gray domain: rest of optic lobe. (C) *Optix-GAL4; UAS-dpp* larvae show more Dm/1/4/12 neurons. (C’) Graphical representation of (C). (D) *Optix-GAL4; UAS-dpp RNAi* larvae show an increase in Dm8 neurons. (D’) Graphical representation of (D). (E) *Optix-GAL4; UAS-brk* larvae have fewer Dm1/4/12 neurons. (E’) Graphical representation of (E). (F) *Optix-GAL4; UAS-brk RNAi* larvae have fewer Dm8 neurons. (F’) Graphical representation of (F). Yellow bracket: Dm8, Pink bracket: Dm1/4/12. Scale bar: 30μm.

### scRNAseq-based lineage tracing identifies a new domain of overlap between Optix and Dpp

Although Brk expression is sufficient to dictate Dm8 *vs.* Dm1/Dm4/Dm12 fates, it is not sufficient to distinguish between Dm1, Dm4 and Dm12 ventrally, or between DRA-Dm8 and Dm11 dorsally. We wondered whether Dpp were also involved in distinguishing between these neural types. As Dpp expression is dynamic during development, we used a *dpp-GAL4*; *UAS-nls-GFP* (Stinger) line, whose expression from the Dpp region perdures in newborn neurons born during larval and early pupal stages. We then FACSed labeled neurons 15 hours after pupation, the earliest time when the scRNAseq transcriptome allows us to define the identity of maturing neurons. We discovered that some neurons appeared to originate from both the Optix and Dpp regions. For example, Dm11, which was shown to be derived from the dorsal Optix region (Figure 2B), was also represented in the Dpp dataset (Figure 6A). Additionally, although Dm1, Dm4 and Dm12 all originated from ventral Optix (Figure 2B), Dm12 was also represented in the Dpp dataset (Dm4 was represented in only one of the two Dpp datasets) (Figure 6A). This suggests that differences in Dpp levels could account for the differences in cell fate between Dm neurons, with high Dpp inducing Dm12 while lower levels would induce Dm4 and then Dm1 (Figure 6A). Staining for Dm1, Dm4 and Dm12 using SoxN and Tj indicated that roughly half of the neurons in the Dm1/4/12 cluster sat within the ventral Dpp region and are most likely Dm12 (Figure 6B-B’’’). In the dorsal Optix domain, Dac and Tj label DRA-Dm8 and Dm11 but some of them were also labeled with *dpp*-driven GFP (Figure 6C-C’’’). As our lineage tracing data suggested that Dm11 are born from both the dorsal Optix and Dpp domains, while DRA-Dm8 cells are only found in datasets from the Optix domain (in very low number in the dOptix dataset, see Table S1)^9^, we conclude that the Dpp^+^/Dac^+^/Tj^+^ cells are likely Dm11 neurons, while the Dpp^-^/Dac^+^/Tj^+^ likely corresponds to both DRA-Dm8 neurons (and possibly Dm11 neurons, Figure 6A, 6C-C’’). This indicates that dorsal Dm11 neurons, like ventral Dm12 neurons, are born from the intersection between the Dpp and Optix regions.

**Figure 6:**
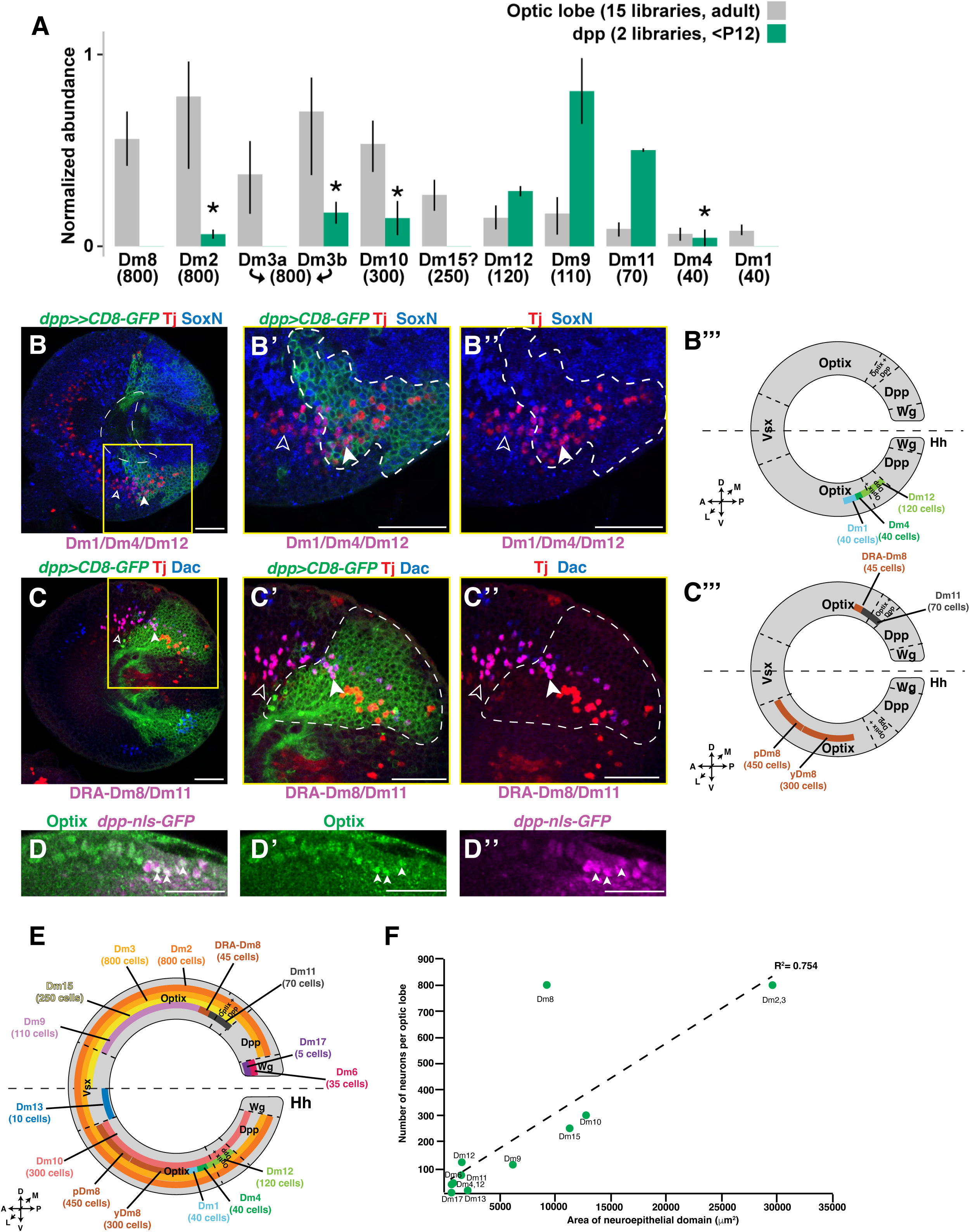
scRNAseq of fluorescently-labeled lineage tracing lines identifies additional spatial patterning required for Dm neuron fate and stoichiometry. (A) *dpp-GAL4; UAS-nls-GFP* lines were FACS sorted and subjected to scRNAseq to identify the spatial region of origin for each cell type. Average normalized abundance of the Dm clusters in datasets produced either in the whole optic lobe or from neurons from the FACSed datasets. The bars represent the minimal and maximal values across all libraries of a dataset. The asterisks indicate when at least 1 library had less than 3 cells of a given cell type, or that the annotations were made with low confidence. Below the graph is the expected stoichiometry of each cell type. (B) SoxN+Tj+ (Dm1/4/12) neurons are produced both inside (carat, likely Dm12) and outside (empty carat, likely Dm1/4) the Dpp expression domain. Dotted line: outline of medulla cortex. (B’-B’’) Inset. Dotted line: region of *dpp>>CD8-GFP* expression. (B’’’) Graphical representation of B-B’’ (See Table S1 for Dm1/4/12 positioning). (C) Some dorsal Dac+Tj+ (DRA-Dm8 and/or Dm11) neurons sit outside the Dpp region (empty carat), while other dorsal Dac+Tj+ (Dm11) neurons sit within the Dpp region. (C’-C’’) Inset. (C’’’) Graphical representation of C-C’’. (D-D’’) Optix protein expression overlaps with *dpp-GAL4*; *UAS-nls-GFP.* Carat: cells with overlapping Dpp/Optix expression. (E) Graphical representation of Dm neuron origins within the OPC (See Table S1). (F) Correlation between neuroepithelial domain size and neuron number. GFP-expressing lines for each spatial factor were imaged and each spatial domain was measured at its widest point; its area was then calculated (Table S2). Each neuron was given a spatial identity, the rationale for which is described in Table S1, and which was then further characterized if the neuron was born from a smaller spatial subdomain (*i.e.,* Dm4 from 1/6 of the ventral Optix domain). The linear plot of OPC domain of origin vs. number of neurons per optic lobe was then plotted for each neuron class (R^2^= 0.754, P=0.0001). For all images, scale bar: 30μm.

Previously, we assumed that Dpp defined an OPC spatial domain distinct from the neighboring Optix domain. However, a *dpp-GAL4* line expressed nuclear-GFP in a pattern that colocalized with endogenous Optix protein at the boundary of the domains (Figure 6D-D’’). This suggests that Dpp and Optix co-expression forms a smaller spatial domain, and that this domain specifies the production of additional neuron types, *e.g.,* Dm11 and Dm12.

We could then make a more accurate map of the spatial origins of Dm neurons (Figure 6E). Neurons that are largest in number, such as Dm2 and Dm3, are born from most of the domains from the main OPC. Neurons that are smaller in number, such as Dm15, are born from two or so subdomains (such as the dorsal Vsx and Optix). Neurons that are in the smallest number are born from single subdomains, some of which have even further spatial subdivisions via Dpp signaling. For example, Dm1 and Dm4 are from the posterior 1/3 of the Optix domain non-overlapping with Dpp, while Dm12 originates from the sub-region that overlaps with Dpp. The numbers of Dm1 and Dm4 neurons together are roughly the same number as Dm12, which is consistent with the relative size of the domains where they are born.

To quantify the relationship between spatial origin and number of each Dm neuron class, we measured the size of each neuroepithelial domain of birth for each Dm neuron at late L3 and fitted a linear model against the final number of each Dm neuron (Table S2). The number of Dm neurons generated appears to be directly proportional to the size of the spatial domain of origin for each neural class (Figure 6F, R^2^= 0.754, P=0.0001). As Dm8s have ∼800 neurons and are born from the anterior 2/3 of the ventral Optix domain and the posterior 1/3 of the dorsal Optix domain, they were a large outlier in the dataset. This exception is discussed below. Despite this exception, our data suggests that spatial patterning regulates the size of the neuroblast pools that generate each type of distal medulla neuron, providing a mechanism that, along with temporal patterning, concurrently regulates cell fate with cell number.

## Discussion

### A gradient of Dpp signaling acts with other spatial factors to promote cell fates in different proportions

Our suggest that multiple spatial patterning pathways intersect to specify neuronal classes with differing stoichiometry (Figure 7). Vsx, Optix and Rx are transcription factors that mutually inhibit each other’s expression to define spatially restricted domains from which neurons of different fates are born^12^. Spalt and Disco act orthogonally and divide the Vsx/Optix/Rx domains to provide dorsal or ventral patterning information^25^. However, these domains are still large, and are not sufficient to explain how neurons with lower abundance are generated. We showed that Dpp signaling regulates Brk expression to generate an additional subdomain. This domain accounts for the distinction between Dm8 and Dm1/4/12 within the Optix domain. However, we were not able to identify the spatial domain that distinguishes p *vs.* yDm8 fate within the Optix/Brk domain. This remains an exciting direction for future study.

**Figure 7:**
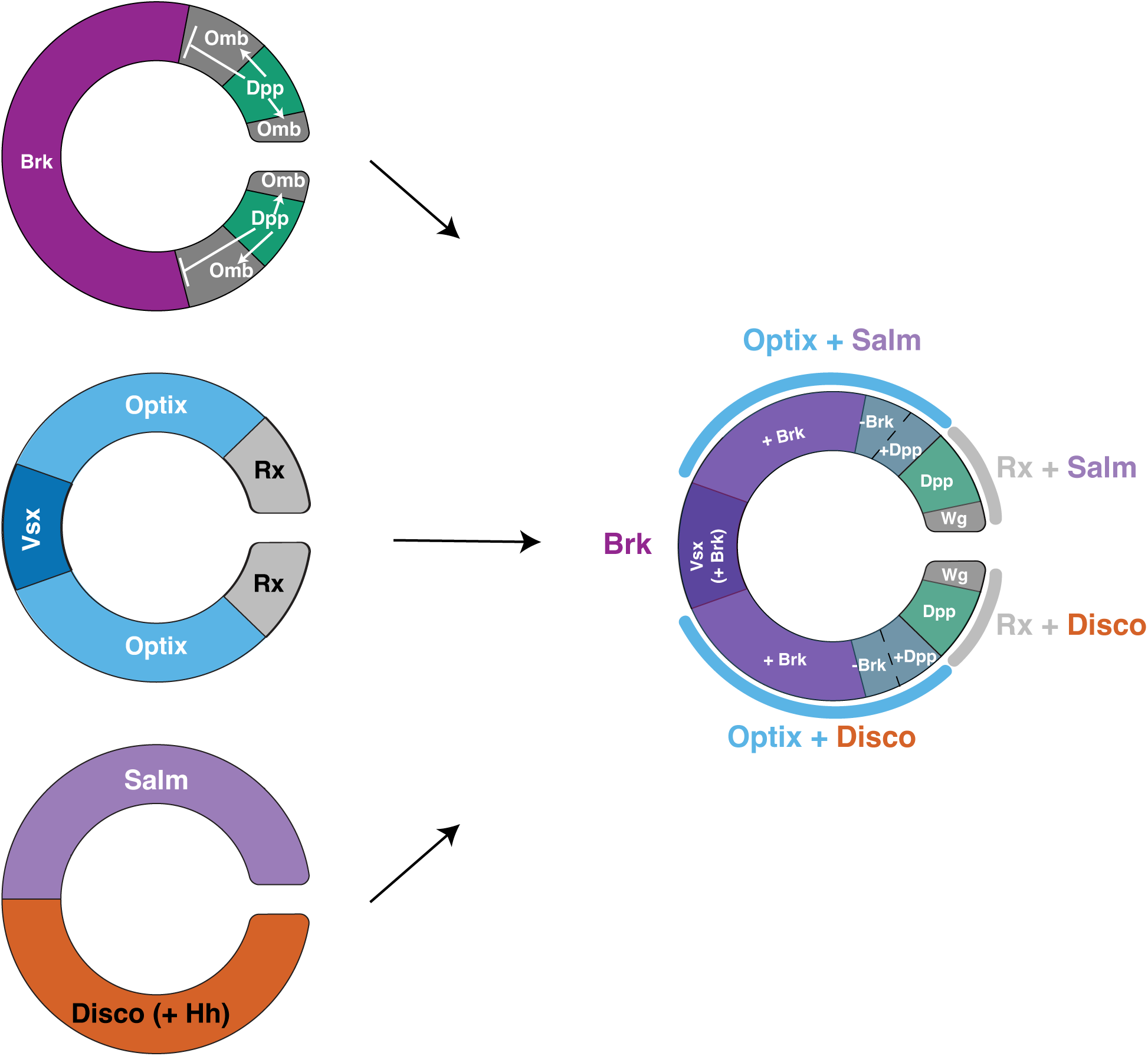
Model: Spatial signaling regulates cell proportions. Three intersecting spatial patterning mechanisms, one using morphogen signaling (Dpp/Brk) and the other two using transcription factor expression (Vsx/Optix/Rx and Salm/Disco+Hh) act together to specify Dm neuron fates in different numbers.

Our previous finding that Dpp defines a subdomain of the Rx region was initially surprising^12^, as Dpp traditionally acts non-autonomously to promote cell fate specification. Our data here confirms that Dpp indeed acts canonically; it activates pMad at a distance, thereby defining the boundary of its negative target, Brinker, and thus splitting the Optix domain into smaller compartments. Our overexpression experiments also suggest that Dpp acts as a morphogen, as its ectopic expression appears to change the distribution of cell types generated during development.

In other systems, the Dpp morphogenetic gradient is converted into discrete domains with sharp boundaries^44^, as Dpp targets contain binding sites of different affinity for its effector, pMad; these interactions can be further refined by cross interactions between Dpp targets^45^. In the OPC, the Brk domain could be determined purely by pMad expression; however, it is also possible that *brk* and *omb* mutually repress each other.

Dpp also appears to play a second role in dividing the Optix^+^ Brk^-^ subdomain in two. The overlap of Dpp and Optix expression distinguishes between cell types both in the ventral and dorsal Optix^+^Brk^-^ domains: Dorsal Dm11 and ventral Dm12 are produced at the region where Dpp overlaps with Optix. This is likely because it is where Dpp signaling is highest, with lower levels of Dpp signaling in the adjacent Optix^+^Brk^-^Dpp^-^ domain. In this case, lower levels of Dpp signaling would specify ventral Dm1/Dm4 and dorsal DRA-Dm8 fates, while the lowest levels of Dpp signaling would allow for increased Brk expression to specify ventral Dm8 fate.

In another region of the *Drosophila* visual system, Dpp and Brk are also used to pattern a second neuroepithelial domain, the Inner Proliferation Center. As with the OPC, Omb is expressed in neurons born from the Dpp region, and Brk is expressed in a neuroepithelial domain that is mutually exclusive to Dpp^45^. These Brk and Dpp domains generate cells that detect global motion in different orientations^46^. pMad is localized to the Dpp domain, and *dpp* RNAi lowers pMad levels and expands the *brk* domain^46–48^. Therefore, Dpp signaling acts across two separate optic anlages to set up domains of expression that promote cell fate specification in the optic lobe.

### Spatial patterning as a mechanism for regulating cell number

Cell proliferation and death are considered to be the major players in the regulation of cell number. Fat/Hippo signaling is the predominant pathway used to promote proliferation in the regulation of organ size. Fat is required to regulate the pace of the transition from OPC neuroepithelial cells to neuroblasts, thereby regulating the switch between symmetric and asymmetric division; in this model, reduction of Fat would delay the movement of the proneural wave, thus expanding the neuroepithelium and the overall size of the optic lobe^49,50^. However, this does not regulate the relative abundance of each neural class. It is still possible that proliferation also regulates cell number in the OPC. For example, a major outlier in the relationship between neuroepithelial domain size and cell number is Dm8, for which 800 neurons are generated from a restricted subdomain of Optix: Dm8 might be born from a longer temporal window or use a transit-amplifying cell to generate additional neurons^51,52^.

Cell death is also used to regulate cellular abundance in the nervous system. Many neuron classes are generated in excess, allowing the appropriate partners to form circuits while culling superfluous cells later during development. In the OPC, it appears that apoptosis is used to fine-tune neuron number, as we have shown that two types of y and pDm8 are made in excess to accommodate the variable number of stochastically made y *vs.* pR7 photoreceptors^9^. The superfluous neurons that do not connect to their partners die by apoptosis. However, cell death does not appear to set the initial number of cells during development.

Our work suggests that the size of the domain of origin in the neuroepithelium for each neuron represents an additional mechanism for regulating cell number. Furthermore, morphogenetic signaling by Dpp can allocate differently sized pools of stem cells to generate different classes of neurons of different numbers during development.

Like the Dm neurons born from the OPC, a large number of neuron classes are born from sheets of neuroepithelia in other systems, many of which are divided into transcriptionally discrete subdomains. For example, the germinal zone that generates the spinal cord and the ventricular zone of the cerebral cortex are also neuroepithelial structures that exhibit expression of spatially restricted transcription factors^53,54^. These systems also use Dpp signaling to set up transcriptional domains. Possibly the best-known example is the vertebrate spinal cord, where the Dpp homolog BMP signals from the vertebrate roofplate to pattern different classes of spinal cord interneurons based on their distance from the initial signal^55,56^. As in the OPC, the output of BMP expression is transduced by the upregulation of distinct classes of transcription factors^57,58^. Therefore, it is possible that this mechanism is used in other contexts to regulate neuronal stoichiometry.

## Acknowledgments

We thank members of the Desplan lab for providing helpful discussions and useful feedback on the manuscript; I. Holguera for advice on Dm transcription factor markers; R. Rajesh and S. Cordoba for advice on figures; R. El-Danaf and K. Kapuralin for advice on Dm6/17 origins; T. Erclik, I. Hariharan, N. Baker, H. Richardson, M. Landgraf, T. Erclik, and D. Ryoo for fly lines; and D. Godt, F. Matsuzaki and R. Mann for antibodies. This work was supported by NIH grant EY019716 and EY10312 to C.D. J.M was supported by NIH fellowships from the National Eye Institute (F32EY028012 and K99EY032269). F.S. was supported by New York University (MacCracken Fellowship and Dean’s Dissertation Ph.D. Fellowship). Y.-C.C. was supported by New York University (MacCracken Fellowship), by a NYSTEM institutional training grant (Contract #C322560GG), and by Scholarship to Study Abroad from the Ministry of Education, Taiwan. This work also benefited from antibodies and *Drosophila* lines generated with the support of the CGSB at NYU in Abu Dhabi.

## Author contributions

J.M. and C.D. conceived the project, analyzed the data and wrote the manuscript. Y.-C.C. performed some immunostains and analyzed scRNAseq data. F.S. performed scRNAseq and analyzed the scRNAseq data. E.K. helped with the apoptosis experiments. J.M. performed all other experiments.

## STAR Methods

### Antibodies

Antibodies:
| Reagent or Resource | Source | Identifier |
| --- | --- | --- |
| Chicken Anti-β-gal | Abcam | Catalog #9361 |
| Rabbit Anti-GFP | Life Technologies | Catalog #11122 |
| Sheep Anti-GFP | Bio-Rad | Catalog #4745-1051 |
| Chicken Anti-GFP | Millipore Sigma | Catalog #06-896 |
| Rabbit Anti-RFP | MBL | Catalog #PM005 |
| Mouse Anti-RFP | MBL | Catalog #M155-3 |
| Rabbit Anti-HA | Cell Signaling Technologies | Catalog #C29F4 |
| Rat Anti-Flag | Novus Biologicals | Catalog #NBP1-06712 |
| Rabbit Anti-Eyeless | Genscript/Desplan Lab | N/A |
| Mouse Anti-Eyeless | DSHB | Anti-Eyeless-s |
| Rabbit Anti-p-Smad3 | Abcam | Catalog #ab52903 |
| Rat Anti-NCadherin | DSHB | DN-Ex #8 |
| Guinea Pig Anti-Traffic Jam | D. Godt | N/A |
| Guinea Pig Anti-Otd | Genscript/Desplan Lab | N/A |
| Rabbit Anti-Dve | F. Matsuzaki | N/A |
| Rabbit Anti-SoxN | Genscript/Desplan Lab | N/A |
| Mouse Anti-Islet/Tup | DSHB | 40.3A4 |
| Anti-Vsx2 | Genscript/Desplan Lab | N/A |
| Mouse Anti-V5-tag:DyLight550 | Bio-Rad | Catalog #MCA1360D550GA |
| Guinea Pig Anti-Lim3 | J. Skeath | N/A |
| Mouse anti-Svp | DSHB | Catalog #5B11 |
| Guinea Pig Anti-Scro | Genscript/Desplan Lab | N/A |
| Mouse Anti-Dac | DSHB | mAbdac-2-3 |
| Guinea Pig Anti-Dac | Genscript/Desplan Lab | N/A |
| Guinea Pig Anti-Otd | Genscript/Desplan Lab | N/A |
| Rat Anti-Otd | Desplan Lab | N/A |
| Rabbit Anti-Toy | Genscript/Desplan Lab | N/A |
| Rat Anti-Toy | Genscript/Desplan Lab | N/A |
| Rabbit Anti-Optix | Genscript/Desplan Lab | N/A |
| Guinea Pig Anti-Brinker | R. Mann | N/A |
| AlexaFluor405 conjugated<br>Goat Anti-HRP | Jackson Immunochemicals | Code #123-475-021 |
| AlexaFluor647 conjugated<br>Goat Anti-HRP | Jackson Immunochemicals | Code #123-605-021 |
| Donkey Anti Rat 405 | Jackson Immunochemicals | Code # 712-475-150 |
| Donkey Anti Rabbit 405 | Jackson Immunochemicals | Code #711-475-152 |
| Donkey Anti Mouse 405 | Jackson Immunochemicals | Code #711-475-150 |
| Donkey Anti Sheep A488 | Jackson Immunochemicals | Code #713-545-147 |
| Donkey Anti Chicken A488 | Jackson Immunochemicals | Code #703-545-155 |
| Donkey Anti Rabbit A488 | Jackson Immunochemicals | Code #711-545-152 |
| Donkey Anti Mouse A488 | Jackson Immunochemicals | Code #715-545-151 |
| Donkey Anti Guinea pig<br>A488 | Jackson Immunochemicals | Code #706-545-148 |
| Donkey Anti Mouse A555 | Thermo Scientific | Catalog #A-31570 |
| Donkey Anti Rabbit A555 | Thermo Scientific | Catalog #A-31572 |
| Donkey Anti Chicken Cy3 | Jackson Immunochemicals | Code # 703-165-155 |
| Donkey Anti Guinea pig Cy3 | Jackson Immunochemicals | Code #706-165-148 |
| Donkey Anti Rat Cy3 | Jackson Immunochemicals | Code #712-165-153 |
| Donkey Anti Guinea pig<br>A647 | Jackson Immunochemicals | Code #706-605-148 |
| Donkey Anti Rabbit A647 | Jackson Immunochemicals | Code #711-605-152 |
| Donkey Anti Mouse A647 | Jackson Immunochemicals | Code #715-605-151 |
| Donkey Anti Rat A647 | Jackson Immunochemicals | Code #712-605-153 |

### Experimental models: Organisms or Strains

| Reagent or Resource | Source | Identifier |
| --- | --- | --- |
| Fly: *D. melanogaster*: *w*,<br>*20xUAS-flpG5::Pest; sp/Cyo*;<br>*TM2/TM6B* | Bloomington | Stock #55807 |
| Fly: *D. melanogaster*:<br>;;*Act5C-FRT-StopFRT-*<br>*lacZ.nls* | Bloomington | Stock #6355 |
| Fly: *D. melanogaster*:<br>*w*;;*13xLexAop2-CD8::GFP* | Bloomington | Stock #32203 |
| Fly: *D. melanogaster*: *w*;<br>*13xLexAop2-CD8::GFP*; | Bloomington | Stock #32205 |
| Fly: *D. melanogaster*:<br>;; *Pxb-GAL4* | Desplan lab, from MI05058 | N/A |
| Fly: *D. melanogaster*: *Optix-*<br>*GAL4<sup>NP2631</sup>* | Kyoto | Stock #104266 |
| Fly: *D. melanogaster*:;; *hh-*<br>*GAL4* | Bloomington | Stock #67046 |
| Fly: *D. melanogaster*: *w1118*;<br>*R22D12-LexA* | Bloomington | Stock #52670 |
| Fly: *D. melanogaster*: *w1118*;<br>*R20D11-LexA* | Bloomington | Stock #52565 |
| Fly: *D. melanogaster*: *w1118*;<br>*R24F10-LexA* | Bloomington | Stock #52696 |
| Fly: *D. melanogaster*: *w<sup>1118</sup>*;<br>*R24F06-LexA* | Bloomington | Stock #52695 |
| Fly: *D. melanogaster*: *w<sup>1118</sup>*;<br>*R42H01-LexA* | Bloomington | Stock #54204 |
| Fly: *D. melanogaster*: *w<sup>1118</sup>*;<br>*R11C05-LexA* | Bloomington | Stock #54608 |
| Fly: *D. melanogaster*: *w<sup>1118</sup>*;<br>*R47G08-LexA* | Bloomington | Stock #52793 |
| Fly: *D. melanogaster*: *w<sup>1118</sup>*;<br>*R18G08-LexA* | Bloomington | Stock #52533 |
| Fly: *D. melanogaster*: *w<sup>1118</sup>*;<br>*R23C03-LexA* | Bloomington | Stock #61523 |
| Fly: *D. melanogaster*: *yw*;<br>*10xUAS-myr-GFP*; | Bloomington | Stock #32198 |
| Fly: *D. melanogaster*:<br>*If/CyO*; *UAS-p35* | Bloomington | Stock #5073 |
| Fly: *D. melanogaster*: ;*Elav*-<br>*GAL4/Cyo-GFP*; *Dr/TM6C* | From Lin and Goodman,<br>1994<sup>59</sup> | N/A |
| Fly: *D. melanogaster*: *w<sup>1118</sup>*;;<br>*R22D12-GAL4* | Bloomington | Stock #48983 |
| Fly: *D. melanogaster*: *w<sup>1118</sup>*;<br>*R26H07-GAL4* | Bloomington | Stock #69854 |
| Fly: *D. melanogaster*: $w^{1118};;$<br>*R20D11-GAL4* | Bloomington | Stock #47891 |
| Fly: *D. melanogaster*: $w^{1118};;$<br>*R24F10-GAL4* | Bloomington | Stock #49090 |
| Fly: *D. melanogaster*: $w^{1118};;$<br>*R24F06-GAL4* | Bloomington | Stock #49087 |
| Fly: *D. melanogaster*: $w^{1118};;$<br>*R42H01-GAL4* | Bloomington | Stock #48150 |
| Fly: *D. melanogaster*: $w^{1118};;$<br>*R11C05-GAL4* | Bloomington | Stock #48291 |
| Fly: *D. melanogaster*: $w^{1118};;$<br>*R47G08-GAL4* | Bloomington | Stock #50328 |
| Fly: *D. melanogaster*: $w^{1118};$<br>*R38A07-GAL4* | Bloomington | Stock #49978 |
| Fly: *D. melanogaster*: $w^{1118};$<br>*R47E05-GAL4* | Bloomington | Stock #50312 |
| Fly: *D. melanogaster*: $w^{1118};;$<br>*R18G08-GAL4* | Bloomington | Stock #47877 |
| Fly: *D. melanogaster*: $w^{1118};;$<br>*R58G11-GAL4* | Bloomington | Stock #39195 |
| Fly: *D. melanogaster*: $w^{1118};;$<br>*VT043152-GAL4* | Vienna Tile | VDRC201362 |
| Fly: *D. melanogaster*: *w*;<br>*UAS-mir(rpr, hid, grim)/CyO; Dr/TM6C* | Iswar Hariharan | N/A |
| Fly: *D. melanogaster*: ;*UAS-Dronc-DN (C318G)*; | Helena Richardson/Nick Baker | N/A |
| Fly: *D. melanogaster*: ;;<br>*UAS-Dronc-RNAi* | Vienna Tile | VDRC100424 |
| Fly: *D. melanogaster*: *w*;;<br>*dronc[L32], FRT80/TM3* | Don Ryoo | N/A |
| Fly: *D. melanogaster*: *tj-GAL4<sup>NP1624</sup>* | Kyoto | Stock #104055 |
| Fly: *D. melanogaster*:<br>;;*Ubi>>stinger, UAS-Flp* | Bloomington | Stock #28282 |
| Fly: *D. melanogaster*:<br>;*ubi>>stinger*; | Bloomington | Stock #32250 |
| Fly: *D. melanogaster*: ;; *dpp-GAL4* | Bloomington | Stock #1553 |
| Fly: *D. melanogaster*:<br>*w;UAS-Stinger*; | Bloomington | Stock #84277 |
| Fly: *D. melanogaster*: ;;*UAS-sf-GFP-nls-PEST-2A-B2ase* | Desplan Lab | N/A |
| Fly: *D. melanogaster*: *yw*,<br>*hsFlp, tub>>GAL4; UAS-EGFP*; | Bloomington | Stock #64767 |
| Fly: *D. melanogaster*: *brk<sup>3sB</sup>-GAL4*;; | Bloomington | Stock #53707 |
| Fly: *D. melanogaster*: *w*;<br>*UAS-dpp/TM3,sb,ser* | Bloomington | Stock #53716 |
| Fly: *D. melanogaster*: *UAS-brk.J* | Bloomington | Stock #36536 |
| Fly: *D. melanogaster*: *w<sup>118</sup>, GMR57C10-FLPL; 10xUAS-MCFO (HA, V5, FLAG) (MCFO3)* | Bloomington | Stock #64087 |
| Fly: *D. melanogaster*: *yv*;;<br>*UAS-dpp RNAi* | Bloomington | Stock #36799 |
| Fly: *D. melanogaster*:<br>*y,sc,sev,v; UAS-brk RNAi/CyO*; | Bloomington | Stock #80438 |
| Fly: *D. melanogaster*:<br>*Ap<sup>rK568</sup>-LacZ/CyO* | Bloomington | Stock #5374 |
| Fly: *D. melanogaster*: *w*;<br>*dvGlut-T2A-QF2, QUAS-LacZ/CyO-DG4; Dr/TM6B (Sb, DG4)* | Matthias Landgraf | N/A |
| Fly: *D. melanogaster*:<br>*w<sup>1118</sup>;salr-T2A-VP16AD/(CyO)* | Ted Erclik | N/A |
| Fly: *D. melanogaster*: *w<sup>1118</sup>, disco-T2A-VP16AD/(FM7c)* | Ted Erclik | N/A |
| Fly: *D. melanogaster*: *w<sup>1118</sup>; optix-T2A-GAL4-DBD/(CyO)*; | Ted Erclik | N/A |

### Chemicals, Peptides and Recombinant Proteins

**Chemicals, Peptides and Recombinant Proteins:**
| Reagent or Resource | Source | Identifier |
| --- | --- | --- |
| SlowFade Gold | Thermo Scientific | Catalog #S36936 |
| Vectashield without DAPI | Vector Lab | H-1000 |

### Software and Algorithms

**Software and Algorithms:**
| Reagent or Resource | Source | Identifier |
| --- | --- | --- |
| Fiji | Schindelin et al., 2012 | https://fiji.sc |
| Imaris | RRID:SCR_007370 | http://www.bitplane.com/imaris/imaris |
| Microsoft Excel | RRID: SCR_016137 | https://www.microsoft.com/en-us/microsoft-365/excel |
| Seurat (v.4.0.1) | RRID: SCR_016341 | https://satijalab.org/seurat/ |
| R (v4.2.0) | RRID: SCR_001905 | https://cran.r-project.org/ |
| GraphPad Prism | RRID: SCR_002798 | https://www.graphpad.com/ |

### Other

**Other:**
| Reagent or Resource | Source | Identifier |
| --- | --- | --- |
| Cell Culture Petri Dish | Thermo Scientific | Catalog #171099 |

### Lead contact for reagent/resource sharing

To obtain information about/resources from this paper, please e-mail the Lead Contact, Claude Desplan.

### Materials availability

Antibodies and fly lines will be distributed from our lab.

### Drosophila strains

Flies were grown on standard cornmeal medium at 25°C 12-hour light/dark cycles (except when specified in listed experiments). Both male and female flies were analyzed for each genotype, but no sex-specific differences were noted. The detailed genotype for each figure is given in Table S3.

### Immunohistochemistry

Fly optic lobes were dissected in PBS and fixed for 15 minutes in 4% formaldehyde (v/w) in 1× PBS at 4°C. After 3 quick washes in 1× PBS, brains were blocked in 1× PBS + 0.4% Triton X-100 (PBST) + 0.5% goat serum for 20 minutes. They were then incubated for 2 days at 4°C in primary antibodies diluted in PBST + 0.5% goat serum. After 3 quick washes in PBST + 2 15-minute washes in PBST, brains were incubated for 1-2 days with secondary antibodies diluted in PBST. After washes, brains were mounted in Slowfade or Vectashield and imaged on a Leica SP8 confocal.

### Antibody dilutions

The following primary antibodies were used for immunofluorescence: rat anti-NCadherin (1:50, DSHB), guinea pig anti-Traffic Jam (1:2500, D. Godt), guinea pig anti-Otd (1:500, Genscript), rabbit anti-Dve (1:1000, F. Matsuzaki), rabbit anti-SoxN (1:250, Genscript), rabbit anti-GFP (1:400, Invitrogen), chicken anti-GFP (1:400, EMD), sheep anti-GFP (1:500, BioRad), mouse anti-RFP (1:400, MBL), mouse Anti-V5-tag:DyLight550 (1:50, BioRad), rat anti-FLAG (1:50, Novus), rabbit anti-HA (1:50, Cell Signaling Technologies), rabbit anti-Eyeless (1:200, Genscript), mouse anti-Eyeless (1:10, DSHB), mouse anti-Islet/Tup (1:100, DSHB), chicken anti-beta-Gal (1:500, Abcam), mouse anti-Svp (1:20, DSHB), guinea pig anti-Scro (1:100, Genscript), mouse anti-Dac (1:20, DSHB, mAbdac2-3), guinea pig anti-Dac (1:1000, Genscript), rabbit anti-Toy (1:300, Genscript), rat anti-Toy(1:50, Genscript), rabbit anti-Optix (1:200, Genscript), rabbit anti-p-Smad3 (1:500, Abcam), rabbit anti-Brk (1:200, R. Mann), rabbit anti-Vsx2 (1:1000, Genscript) guinea pig anti-Lim3 (1:500, J. Skeath), and AlexaFluor405 conjugated Goat Anti-HRP (Jackson Immunochemicals, 1:100). Secondary antibodies are from Invitrogen and used at 1:200.

### Quantification and Statistical Analyses

Male and female larvae at similar stages were selected randomly from the fly vials for all experiments. Blinding across different genotypes was not performed, as the genotype can be distinguished by the experimenter. All immunohistochemistry experiments were performed in at least 3 different biological replicates (brains of different flies) for each genotype, which is in line with experiments from other scientists in the field. Apoptosis inhibition led to aberrant neuron targeting. To distinguish between off-target labeling and aberrantly targeting neurons, we only counted cells that were the same size as normally targeting neurons, and we only counted cells that were closest in proximity to normally targeting neurons. Quantification of Dm1/4/12 vs. Dm8/11 cell number was calculated using the “Spots” function on Imaris; only neurons within the Z-plane of the medulla were counted. For all T-tests, a two-tailed T test was performed. T-tests, Chi-square analysis and linear regression were all performed using GraphPad Prism. Averages and standard deviation/standard error of the mean were calculated using Microsoft Excel.

### Image Acquisition and Processing

All images were captured on a Leica SP8 Confocal Microscope. Images were analyzed in FIJI (ImageJ) and Imaris.

### scRNAseq

The raw data was obtained from (Simon et al., in preparation). Briefly, we acquired the data by producing lines expressing a nuclear GFP in neurons from the *Optix*, *vOptix*, *hh*, *dpp* and *pxb* regions of the mOPC, sorting the labelled cells by FACS, and performing single-cell mRNA sequencing on the obtained cell suspension.

The raw data was then analyzed using Seurat 4.0.1. For each library, a Seurat Object was created with all genes expressed at least in 3 cells, and all cells expressing at least 200 genes. The objects were then filtered by keeping all cells below a percent of mitochondrial genes, below a specific number of UMIs, and above a number of genes, based on the distribution of these parameters (plotted in Simon et al., in preparation). The threshold chosen were identical for all libraries acquired a given day with flies of a given genotype, but were different otherwise. For the 3 Optix libraries the thresholds were 7/17000/800 (percent of mitochondrial genes, number of UMIs, number of genes), for the 2 vOptix libraries 10/10000/500, for the 2 dOptix libraries 5/20000/1000, for the 2 hh libraries 10/20000/700, for the 2 dpp libraries 5/20000/900, for the pxb library 5/30000/1300. After filtering, the number of cells in each library was 5532 (Optix 1), 6786 (Optix 2), 6718 (Optix 3), 5175 (vOptix 1), 5421 (vOptix 2), 6182 (dOptix1), 6120 (dOptix2), 4433 (hh 1), 4749 (hh 2), 6127 (dpp 1), 6735 (dpp 2), 4988 (pxb). For each library we then ran NormalizeData, FindVariableFeatures and ScaleData with default parameters, as well as RunPCA, RunTSNE and RunUMAP with defaults parameters and a dimensionality of 150.

Dimensionality reductions were run purely for visualization purposes: these were not used to annotate the dataset, and neither did we perform any clustering. Instead, we used the normalized expression of marker genes and the neural network classifier built and presented in^30^ to assign each cell of each library to its corresponding cluster in our published single-cell atlas (metadata fields “NN_Cluster_Number”), and to give this assignment a confidence score between 0 and 1 (metadata field “Confidence_NN_Cluster_Number”). For all figures, we replaced cluster numbers by cluster annotations (metadata fields “Annotation”) previously published in ^30^, except for Dm15 cluster that we newly identified (Fig. 2B).

For each dataset, if at least 80% of the cells of a given class (group of cells with the same annotation) were annotated with a confidence score strictly below 0.5, the class was flagged as “low confidence”.

Then, we normalized class (group of cells with the same annotation) abundances to allow us to compare them between libraries. To do so, we used the abundances of neuronal clusters produced in the whole mOPC, since they represent a constant between mOPC regions. In each library of our spatial origin datasets, as well as our single cell atlas, we therefore divided the abundance of each class by the abundance of neuronal clusters T1, Mi1, Tm1, Tm2, Tm4, and Tm6, which were produced in all mOPC regions as determined in Simon et al., in preparation.

We then averaged these normalized abundances and plotted them, as well as the minimal and maximal abundances, on Figure 2B and Figure 6A. Finally, for each dataset, we flagged classes containing either 3 cells or less, or for which at least 80% of the cells were annotated with a confidence score strictly below 0.5.

### Correlation of spatial subdomain with Dm neuron number

GFP-expressing lines for each spatial factor were imaged and each spatial domain was measured at its widest point; its area was then calculated (using the lasso tool in ImageJ/FIJI). Each neuron was given a spatial identity, which was then further characterized if the neuron was born from a smaller spatial subdomain (*i.e.,* Dm4 from 1/6 of the ventral Optix domain). The linear plot of OPC subdomain of origin vs. number of neurons per optic lobe was then plotted for each neuron class. A linear regression was performed, and R-square and P value were calculated using Microsoft Excel and GraphPad Prism.

## Supplementary Table S3: Genotypes

Figure 1:

**1B:** R57C10-FlpL/+;; 10XUAS-FRT>STOP>FRT-myr::smGFP-HA,

10XUASFRT>STOP>FRT-myr::smGFP-V5-THS-10XUAS-FRT>STOP>FRT-myr::smGFP-FLAG/R47G08-GAL4 (Nern et al., 2015)

**1C:** R57C10-FlpL/+;; 10XUAS-FRT>STOP>FRT-myr::smGFP-HA,

10XUASFRT>STOP>FRT-myr::smGFP-V5-THS-10XUAS-FRT>STOP>FRT-myr::smGFP-FLAG/R24F06-GAL4 (Nern et al., 2015)

**1D:** R57C10-FlpL/+;; 10XUAS-FRT>STOP>FRT-myr::smGFP-HA,

10XUASFRT>STOP>FRT-myr::smGFP-V5-THS-10XUAS-FRT>STOP>FRT-myr::smGFP-FLAG/R11C05-GAL4 (Nern et al., 2015)

Figure 2:

2C/C’: w, 20xUAS-flpG5::Pest; Act>>nuLacZ/R22D10-LexA; Pxb-GAL4/LexAOp-CD8-GFP

**2D/D’:** w, 20xUAS-flpG5::Pest; Optix-GAL4, LexAOp-CD8-GFP/R22D10-LexA;

Act>>nuLacZ/LexAOp-CD8-GFP

**2E/E’:** w, 20xUAS-flpG5::Pest; Act>>nuLacZ/R22D10-LexA; hh-GAL4. LexAOp-CD8-GFP/LexAOp-CD8-GFP

**2G/G’:** w, 20xUAS-flpG5::Pest; Act>>nuLacZ/R18G08-LexA; Pxb-GAL4/LexAOp-CD8-GFP

**2H/H’:** w, 20xUAS-flpG5::Pest; Optix-GAL4, LexAOp-CD8-GFP/R18G08-LexA;

Act>>nuLacZ/+

**2I/I’:** w, 20xUAS-flpG5::Pest; Act>>nuLacZ/R18G08-LexA; hh-GAL4. LexAOp-CD8-GFP/+

**2K/K’:** w, 20xUAS-flpG5::Pest; Act>>nuLacZ/R24F10-LexA; Pxb-GAL4/LexAOp-CD8-GFP

**2L/L’:** w, 20xUAS-flpG5::Pest; Optix-GAL4, LexAOp-CD8-GFP/R24F10-LexA;

Act>>nuLacZ/+

**2M/M’:** w, 20xUAS-flpG5::Pest; Act>>nuLacZ/R24F10-LexA; hh-GAL4. LexAOp-CD8-GFP/+

Figure 3:

**3A:** w/yw; R22D12-LexA/Elav-GAL4; LexAOp-CD8-GFP/UAS-p35 (used males from cross, Elav-GAL4; UAS-p35 line can not be maintained)

w/yw; R20D11-LexA/Elav-GAL4; LexAOp-CD8-GFP/UAS-p35 w/yw; R24F10-LexA/Elav-GAL4; LexAOp-CD8-GFP/UAS-p35 w/yw; R38H06-LexA/Elav-GAL4; LexAOp-CD8-GFP/UAS-p35 w/yw; R42H01-LexA/Elav-GAL4; LexAOp-CD8-GFP/UAS-p35

w/yw; R11C05LexA/Elav-GAL4; LexAOp-CD8-GFP/UAS-p35 w/yw; R47G08-LexA/Elav-GAL4; LexAOp-CD8-GFP/UAS-p35 w/yw; R18G08-LexA/Elav-GAL4; LexAOp-CD8-GFP/UAS-p35 w/yw; R23C03-LexA/Elav-GAL4; LexAOp-CD8-GFP/UAS-p35 **3B:** yw; UAS-CD8-GFP; R24F10-GAL4

yw; UAS-CD8-GFP/+; R24F10-GAL4/UAS-RHG-miRNA yw; UAS-CD8-GFP/+; R24F10-GAL4/UAS-Dronc-DN yw; UAS-CD8-GFP/+; R24F10-GAL4/UAS-Dronc-RNAi yw; R24F10-LexA, LexAOp-CD8-GFP; TM2/TM6

yw; R24F10-LexA, LexAOp-CD8-GFP; *dronc[L32]*

yw; R24F10-LexA, LexAOp-CD8-GFP; tj-GAL4/TM2

yw; R24F10-LexA, LexAOp-CD8-GFP; tj-GAL4/UAS-p35 yw; R24F10-LexA, LexAOp-CD8-GFP/Elav-GAL4; +/TM2

yw; R24F10-LexA, LexAOp-CD8-GFP/ Elav-GAL4; +/UAS-p35

Figure 4:

**4B/B’:** yw;;

**4C:** brk-GAL4; Sp/CyO; 2AB∼UAS-sf-GFP-nls-PEST

**4D:** w/brk[X47]; Optix-Gal4/+; UAS-Stinger

**4E: ;;** tkv-GAL4/ 2AB∼UAS-sf-GFP-nls-PEST

**4G:** yw, hsFlp, tub>>Gal4; UAS-EGFP; UAS-mCherry RNAi

**4H:** w/brk[X47]; Optix-Gal4/+; UAS-Stinger

**4I:** omb-GAL4, UAS-EGFP; ;

**4J** brk-GAL4; Sp/CyO; 2AB∼UAS-sf-GFP-nls-PEST

**4L/L:** hsflp, tub>>GAL4/+; UAS-EGFP/brk-LacZ; +/UAS-dpp

Figure 5:

**5B:** yw;;

**5C:** yw; Optix-GAL4/+; UAS-dpp/+

**5D:** yw; Optix-GAL4/+; UAS-dpp RNAi/+

**5E:** yw; Optix-GAL4/+; UAS-brk-J./+

**5F**: yw; Optix-GAL4/UAS-brk-RNAi; +/+

Figure 6:

**6B-B’’, 6C-C’’:** w, 20xUAS-flpG5::PEST; Gal80ts; Act>y+>lexA, lexAop-myr-GFP/dpp-Gal4 (48hr ts)

**6D-D’’:** ;; 2AB∼UAS-sf-GFP-nls-PEST/dpp-GAL4

Supplementary Figure 1:

**S1A:** w; Rx-T2A-GAL4; UAS-CD8-GFP

**S1B’:** w, 20xUAS-flpG5::Pest; Rx-T2A-GAL4, tubGal80ts; ubi>>Stinger (grown at 25°C) **S1D/D’:** w, 20xUAS-flpG5::Pest; Act>>nuLacZ/R22D12-LexA; Pxb-GAL4/LexAOp-CD8-GFP **S1E/E’:** w, 20xUAS-flpG5::Pest; Optix-GAL4, LexAOp-CD8-GFP/R22D12-LexA;

Act>>nuLacZ/+

**S1F/F’:** w, 20xUAS-flpG5::Pest; Act>>nuLacZ/R22D12-LexA; hh-GAL4. LexAOp-CD8-GFP/LexAOp-CD8-GFP

**S1H/H’:** w, 20xUAS-flpG5::Pest; Act>>nuLacZ/R38H06-LexA; Pxb-GAL4/LexAOp-CD8-GFP

**S1I/I’:** w, 20xUAS-flpG5::Pest; Optix-GAL4, LexAOp-CD8-GFP/R38H06-LexA;

Act>>nuLacZ/+

**S1J/J’:** w, 20xUAS-flpG5::Pest; Act>>nuLacZ/R38H06-LexA; hh-GAL4. LexAOp-CD8-GFP/LexAOp-CD8-GFP

**S1L/L’:** w, 20xUAS-flpG5::Pest; Act>>nuLacZ/R24H06-LexA; Pxb-GAL4/LexAOp-CD8-GFP

**S1M/M’:** w, 20xUAS-flpG5::Pest; Optix-GAL4, LexAOp-CD8-GFP/R24F06-LexA;

Act>>nuLacZ/+

**S1N/N’:** w, 20xUAS-flpG5::Pest; Act>>nuLacZ/R24F06-LexA; hh-GAL4. LexAOp-CD8-GFP/LexAOp-CD8-GFP

**S1P/P’:** w, 20xUAS-flpG5::Pest; Act>>nuLacZ/R42H01-LexA; Pxb-GAL4/LexAOp-CD8-GFP

**S1Q/Q’:** w, 20xUAS-flpG5::Pest; Optix-GAL4, LexAOp-CD8-GFP/R42H01-LexA;

Act>>nuLacZ/+

**S1R/R’:** w, 20xUAS-flpG5::Pest; Act>>nuLacZ/R42H01-LexA; hh-GAL4. LexAOp-CD8-GFP/LexAOp-CD8-GFP

**S1T/T’:** w, 20xUAS-flpG5::Pest; Act>>nuLacZ/R11C05-LexA; Pxb-GAL4/LexAOp-CD8-GFP

**S1U/U’:** w, 20xUAS-flpG5::Pest; Optix-GAL4, LexAOp-CD8-GFP/R11C05-LexA;

Act>>nuLacZ/+

**S1V/V’:** w, 20xUAS-flpG5::Pest; Act>>nuLacZ/R11C05-LexA; hh-GAL4. LexAOp-CD8-GFP/LexAOp-CD8-GFP

**S1X/X’:** w, 20xUAS-flpG5::Pest; Act>>nuLacZ/R47G08-LexA; Pxb-GAL4/LexAOp-CD8-GFP

**S1Y/Y’:** w, 20xUAS-flpG5::Pest; Optix-GAL4, LexAOp-CD8-GFP/R47G08-LexA;

Act>>nuLacZ/+

**S1Z/Z’:** w, 20xUAS-flpG5::Pest; Act>>nuLacZ/R47G08-LexA; hh-GAL4. LexAOp-CD8-GFP/LexAOp-CD8-GFP

**S1BB/BB’:** w, 20xUAS-flpG5::Pest; Act>>nuLacZ/R38A07-LexA; Pxb-GAL4/LexAOp-CD8-GFP

**S1CC/CC’:** w, 20xUAS-flpG5::Pest; Optix-GAL4, LexAOp-CD8-GFP/R38A07-LexA;

Act>>nuLacZ/LexAOp-CD8-GFP

**S1DD/DD’:** w, 20xUAS-flpG5::Pest; Act>>nuLacZ/R38A07-LexA; hh-GAL4. LexAOp-CD8-GFP/LexAOp-CD8-GFP

**S1FF/FF’:** w, 20xUAS-flpG5::Pest; Act>>nuLacZ/R23C03-LexA; hh-GAL4. LexAOp-CD8-GFP/LexAOp-CD8-GFP

**S1GG/GG’:** w, 20xUAS-flpG5::Pest; Optix-GAL4, LexAOp-CD8-GFP/R23C03-LexA;

Act>>nuLacZ/LexAOp-CD8-GFP

**S1HH/HH’:** w, 20xUAS-flpG5::Pest; Act>>nuLacZ/23C03-LexA; hh-GAL4. LexAOp-CD8-GFP/LexAOp-CD8-GFP

Supplementary Figure 3:

**S3A-B’:** yw; UAS-CD8-GFP; VT043152-GAL4 **S3C-C’:** yw; R38H06-LexA; LexAOp-CD8-GFP **S3D-E’:** yw; UAS-CD8-GFP; R38A07-GAL4 **S3F-F’:** yw; UAS-CD8-GFP; R47E05-GAL4 **S3G-G’:** yw; UAS-CD8-GFP; R58G11-GAL4

Supplementary Figure 4:

**S4A:** yw; vGlut-QF, QUAS-nuLacZ/UAS-CD8-GFP; UAS-CD8-GFP/R18G08-GAL4

**S4B:** Ap-LacZ/UAS-CD8GFP; R18G08-GAL4/+

**S4C-D’:** UAS-CD8-GFP; R18G08-GAL4

Supplementary Figure 5:

**S5A/A’:** yw; UAS-stinger; dpp-GAL4

**S5B/B’:** 20XUAS-Flp; Optix-GAL4, LexAOp-CD8GFP; Act>>nuLacZ

**S5D/D’:** yw;;

Supplementary Figure 6:

**S6A-B’’, S6E-E’:** 20XUAS-Flp; Optix-GAL4, tubGal80ts; ubi>>stinger

**S6C-D’:** brk-T2A-GAL4; Sp/CyO; 2AB∼UAS-sf-GFP-nls-PEST

## Supplementary Figure Legends

**Supplementary Figure S1.**
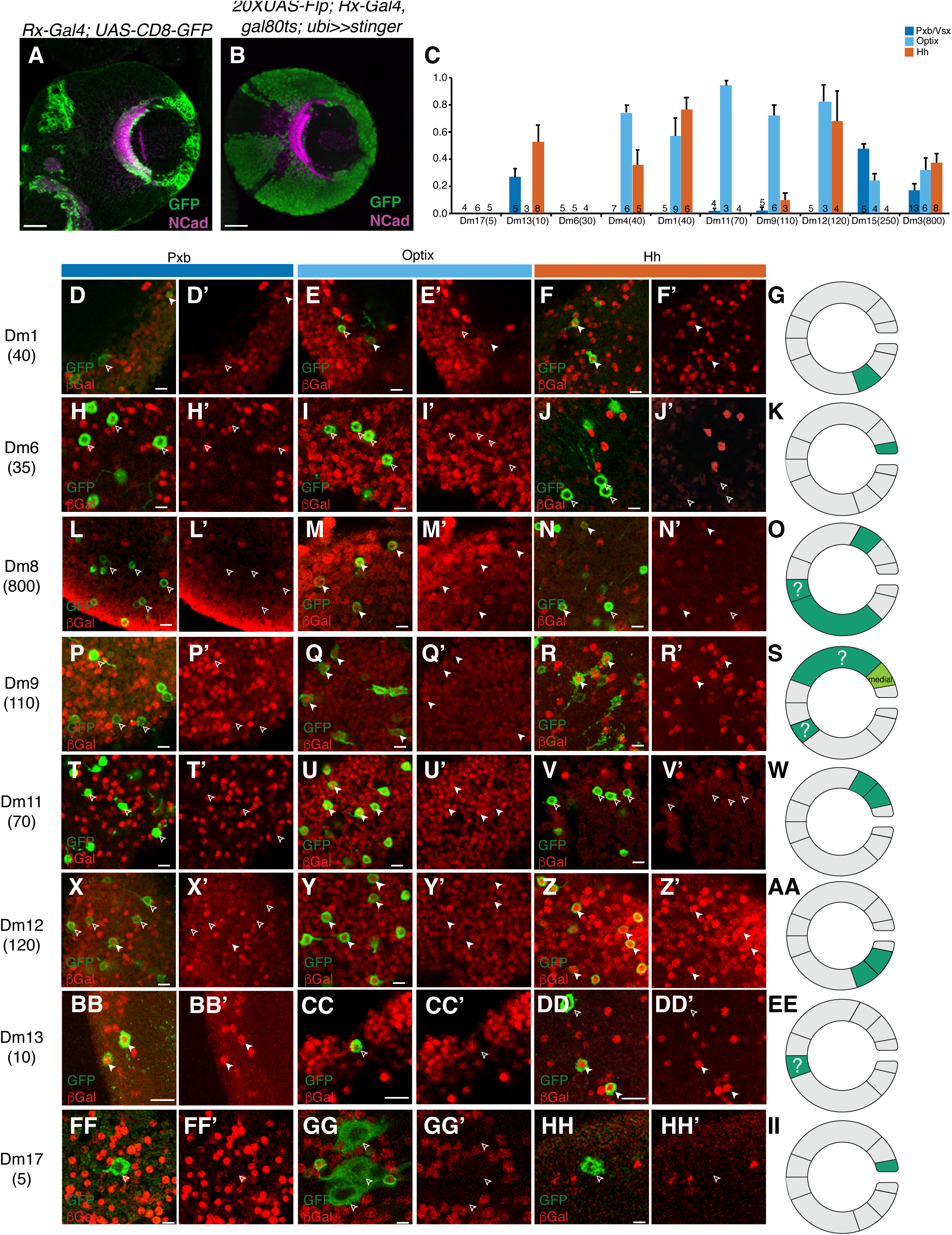
Spatial domain fate mapping of distal medulla neurons. (A) *Rx-GAL4; UAS-CD8-GFP* is expressed in the posterior-most segment in the OPC. (B) An Rx lineage tracing line (*20XUAS-Flp; Rx-GAL4, tubGal80ts; ubi>>stinger*, raised at 25°C) labels all neurons born from the Rx and Optix domains of the OPC, indicating that Rx and Optix expression initially overlaps, but is later refined. Additionally, this expression is visible even when the line is not raised at the permissive temperature, indicating that *Rx-T2A-GAL4* cannot be used for lineage tracing. (C) Quantification of *act>>nuLacZ* lineage tracing data. Scale bar: standard deviation. (D-II) Representative images of lineage trace lines crossed to GFP reporters for each Dm neuron. Open carat: lack of expression; Closed carat: presence of expression. Scale bar = 5μm. (G, K, O, S, W, AA, II) Graphical representation of Dm origins based on lineage tracing data, as well as other experiments (see Table S1).

**Supplementary Figure S2:**
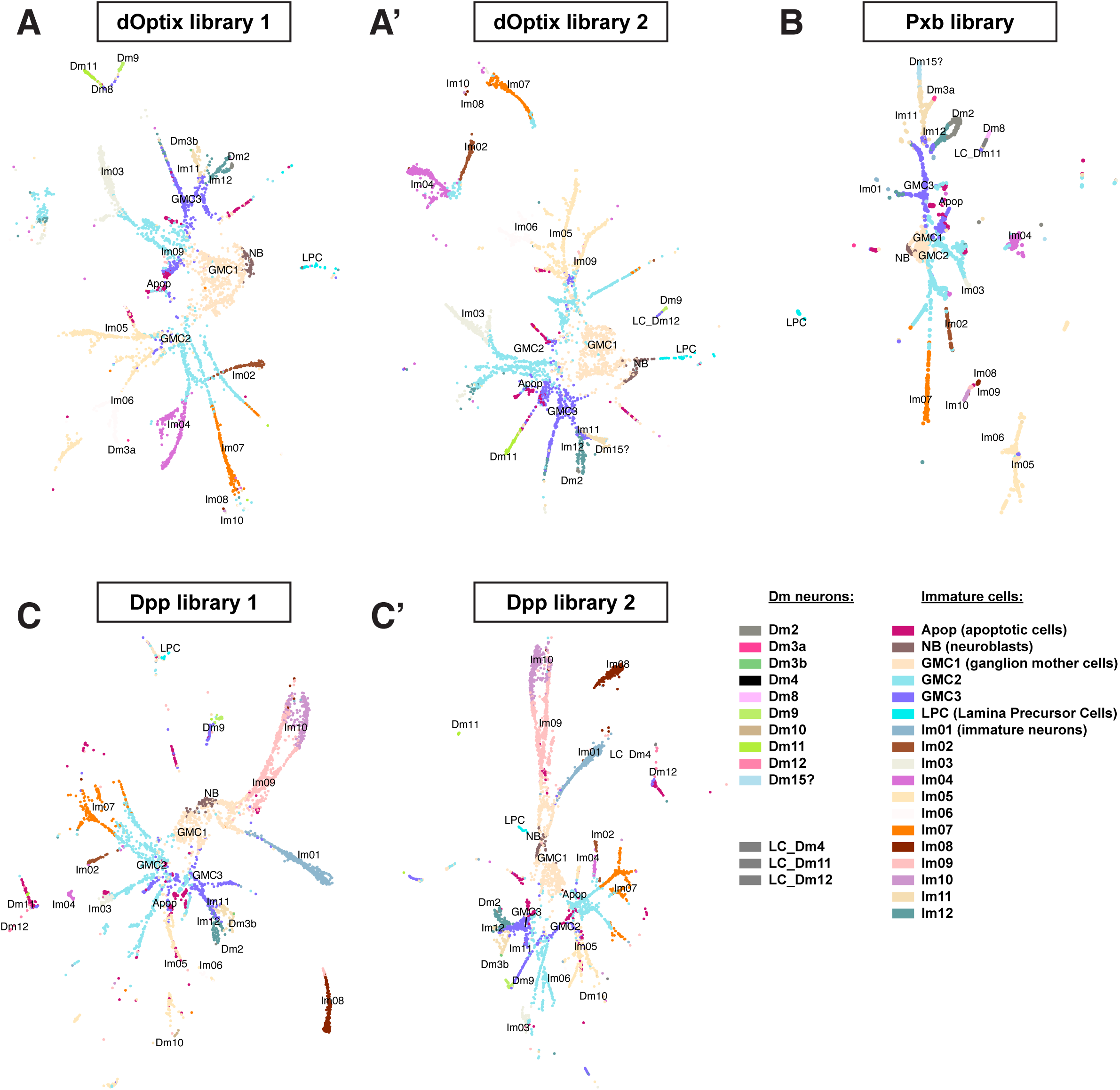
UMAP trajectories of FACSed lineage tracing lines suggest that immature Dm neurons are born from specific spatial subdomains. (A, A’) UMAP plots suggest that immature Dm2, Dm3b (and possibly Dm3a), Dm9, Dm11 and Dm15 neurons are found among the dOptix lineage trace data. (B) UMAP plots suggest that immature Dm2, Dm3a, and Dm15 neurons are found among the Pxb lineage trace data. (C, C’) UMAP plots suggest that immature Dm3b, Dm10, Dm11 and Dm12 neurons are found among the Dpp lineage trace data. For neuronal types annotated “LC_”, the annotation assigned by our neural network was of lower confidence (Methods). This often has a biological explanation: for instance, Dm1/4/12 are very similar and therefore more difficult to annotate by our neural network.

**Supplementary Figure S3:**
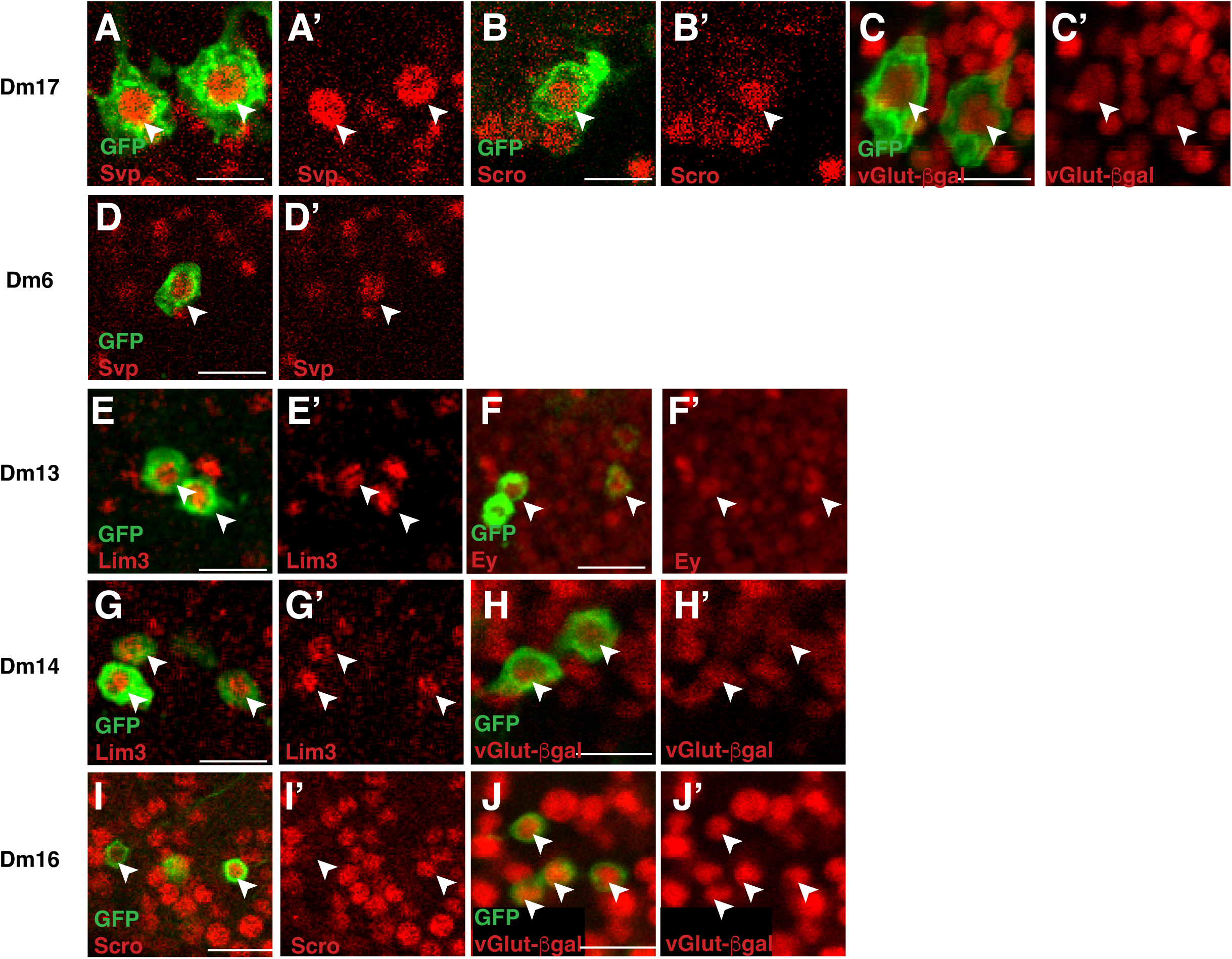
Dm neurons express specific transcription factor markers. (A-J’) Adult *Drosophila* expressing GFP reporters for neurons of interest were immunostained for cell type-specific markers; each Dm subtype expresses distinct sets of transcription factors; carat: cell with transcription factor expression, Scale bar: 10μm. (A-C’) Dm17 expresses Svp, Scro and *vGlut-nuLacZ*. (D-D’) Dm6 expresses Svp. (E-F’) Dm13 expresses Lim3 and Ey. (G-H’) Dm14 expresses Lim3 and *vGlut-nuLacZ*. (I-J’) Dm16 expresses Scro and *vGlut-nuLacZ*.

**Supplementary Figure S4:**
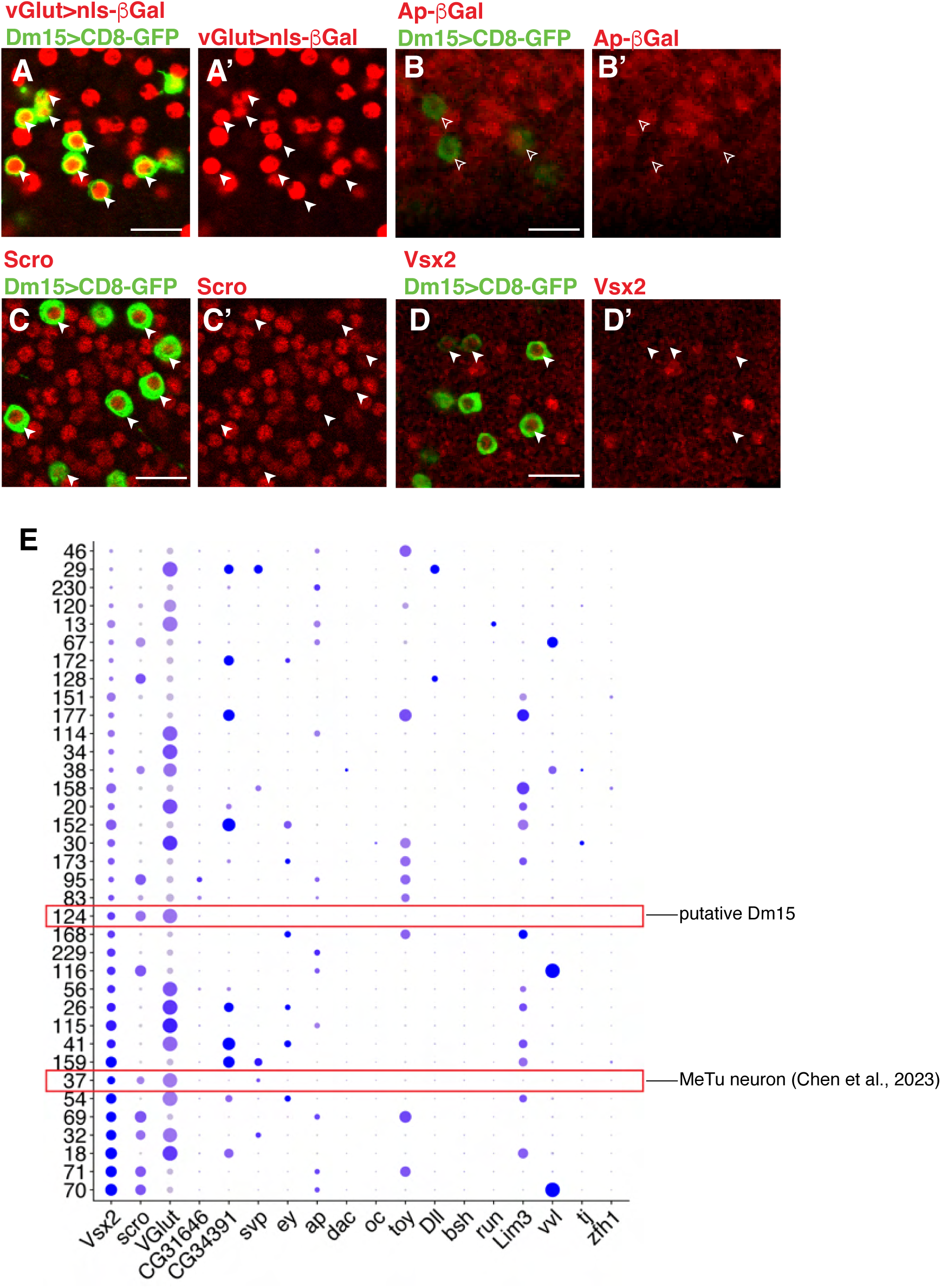
Dm15 is cluster 124 in our scRNAseq dataset. For all images, scale bar: 10μm. Carat: cell that expresses marker of interest. Empty carat: cell that does not express marker of interest. GFP-labeled Dm15 neurons express vGlut (A-A’) but not Apterous (B-B’) in adult animals. They also express Scro (C-C’), and Vsx2 (D-D’). (E) Dot plot of mixture modeling of adult scRNAseq data set (Özel et al., 2021) suggests that Cluster 124 expresses many of the same markers as Dm15. Cluster 37 expresses similar markers but also expresses Svp. A split-GAL4 line that specifically labels cluster 37 has identified it as a MeTu neuron^60^. Red boxes: clusters expressing markers, x-axis: gene name; y axis: cluster number.

**Supplementary Figure S5.**
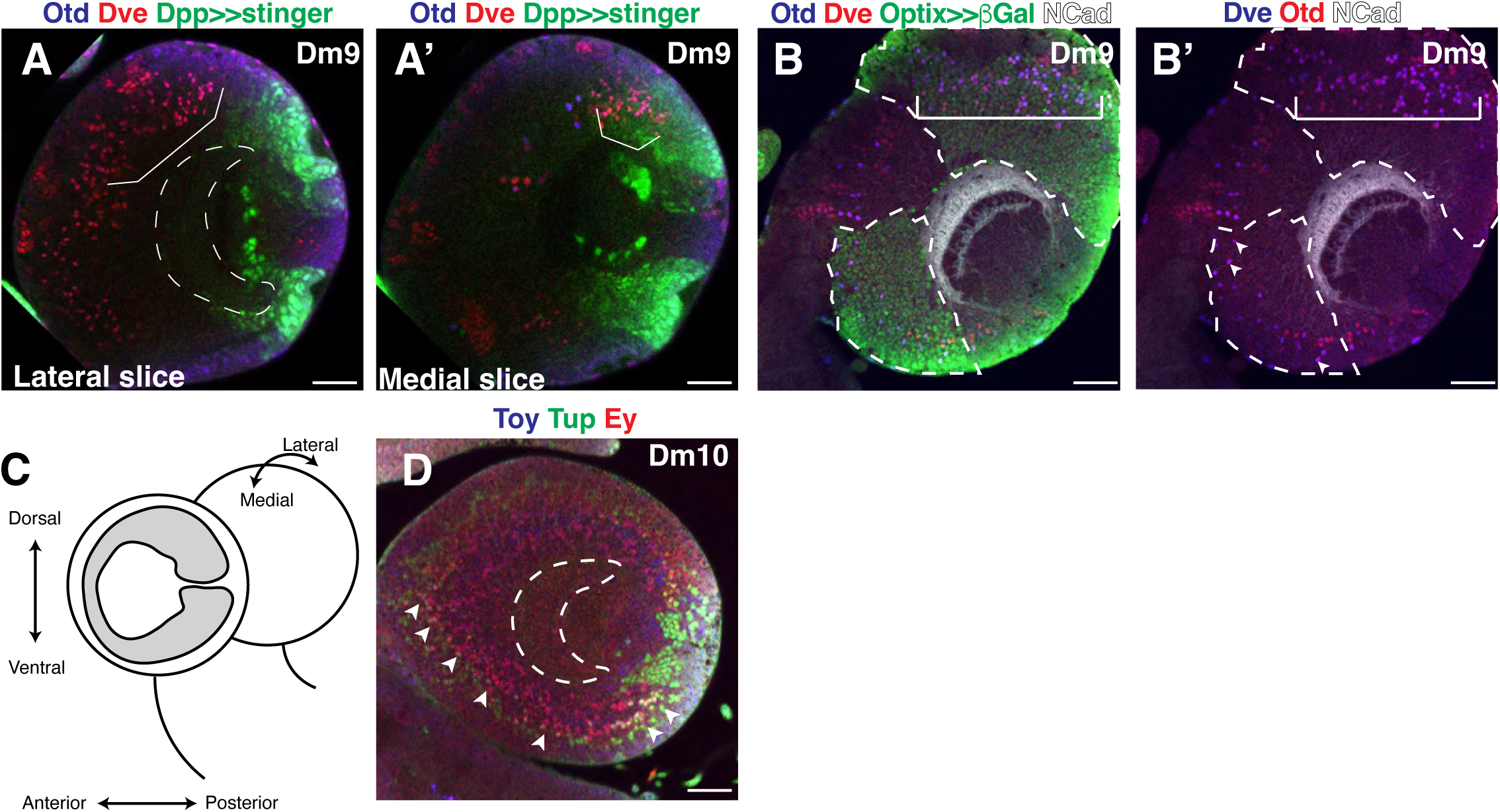
Immunostains against Dm-specific transcription factors place each neuron type in a specific OPC subdomain. (A) Otd^+^Dve^+^ Dm9 neurons are mainly located in the dorsal medulla and are not located within the *UAS-Stinger; dpp-GAL4* domain in the lateral medulla. Dotted line: medulla neuropil position; bracket: Dm9 neurons. (A’) In the medial medulla, Otd^+^Dve^+^ neurons are situated within the Dpp domain. Bracket: Dm9 neurons. (B) Otd^+^Dve^+^ neurons sit within the anterior 2/3 of the dorsal Optix region in the lateral medulla. Bracket: dorsal Dm9 neurons. (B’) Image B without GFP; Carat: smaller population of Otd^+^Dve^+^ Dm9(?) neurons in anterior 1/3 of ventral Optix domain, with a handful of cells in the posterior 1/3 of the ventral Optix domain. (C) A map of the different axes of the optic lobe. Note the orientation of medial to lateral refers to the optic lobe in the background, not the foreground (Adapted from Gold and Brand 2014^24^). (D) Dm10 neurons are situated throughout the entirety of the ventral Optix + Dpp domains. Carat: Toy^+^Tup^+^Ey^+^ Dm10 neurons (white). Scale bar: 30μm.

**Supplementary Figure S6.**
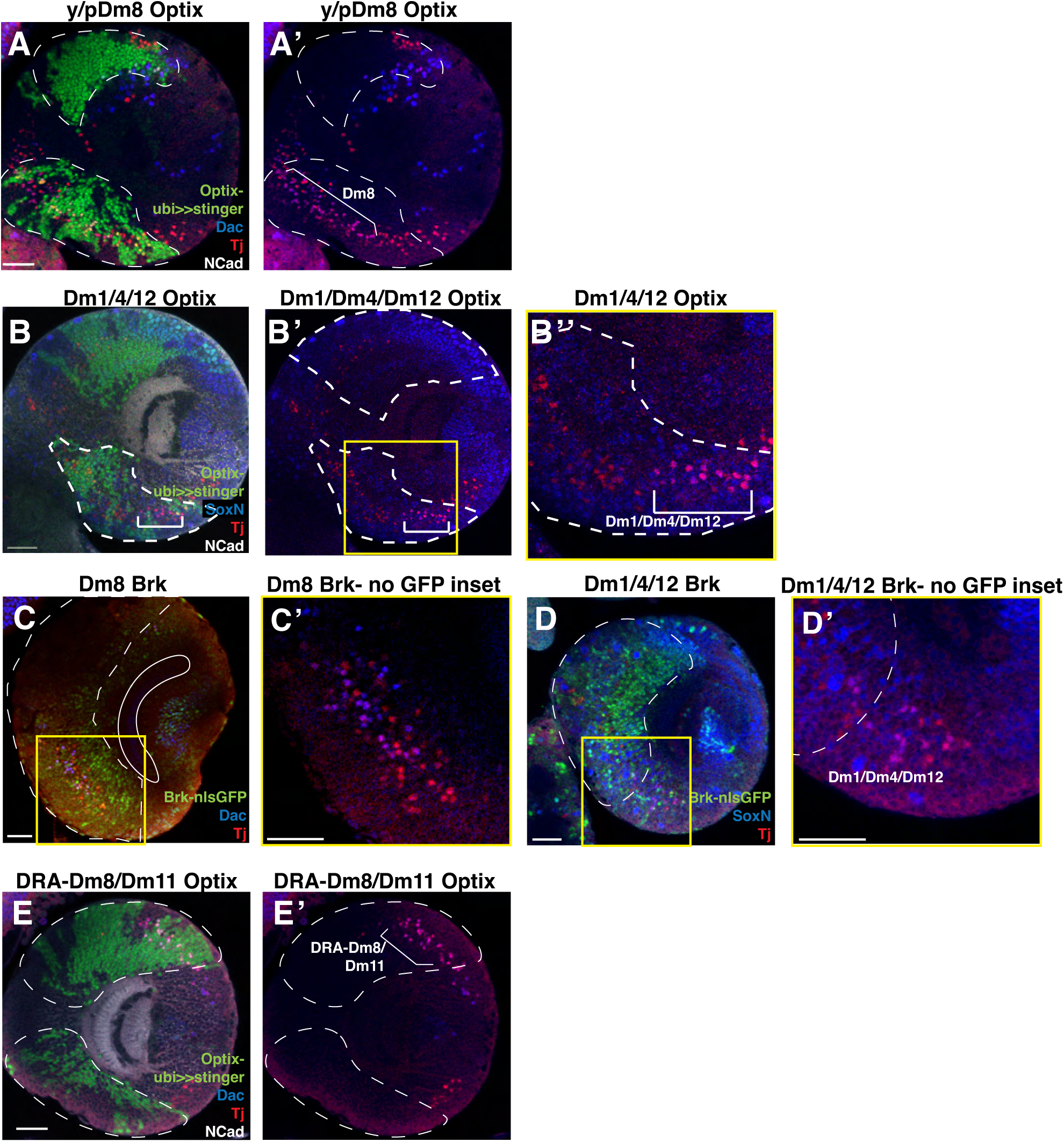
Some Dm neurons are born from Optix subdomains. Brk expression separates specific Dm subtypes. (A-E’) Dotted line: spatial domain boundary. Bracket: neurons of interest. Yellow square: inset. Scale bar: 30μm. (A’-A’) Dac^+^Tj^+^ Dm8 neurons sit within the anterior 2/3 of the ventral Optix domain. Dac^+^Tj^+^ neurons from the Vsx domain could also be Dm8 neurons. (B-B’’) SoxN^+^Tj^+^ Dm1, Dm4 and Dm12 neurons sit within the posterior 1/3 of the ventral Optix domain. (B’) Image without GFP, (B’’) inset of (B’). (C) Solid crescent: outline of medulla neuropil. (C) Dac^+^Tj^+^ neurons sit entirely within the Brk domain. Dotted line= *brk-GAL4;;UAS-nls-sfGFP* expression domain. (C’) Inset of (C) without GFP. (D) Most Dm1, Dm4 and Dm12 neurons sit outside the *brk* domain. A handful sit within the domain. (D’) Inset of (D) without GFP. (E-E’) Dac^+^Tj^+^ DRA Dm8/Dm11 neurons reside within the posterior 1/3 of the dorsal Optix domain.

