## Supplementary Table S1 for "Spatial patterning regulates neuron numbers in the *Drosophila* visual system"

| **Neuron name (cell number)** | **Lineage tracing data** | **scRNAseq data** | **Immunofluorescence data** | **Notes** | **Predicted origin** |
| --- | --- | --- | --- | --- | --- |
| Dm1 (40) | Pxb-, Optix+, Hh+ | Optix+, Hh+, vOptix+, Dpp- | SoxN+Tj+ (which corresponds to Dm1, 4 and 12) = posterior 1/3 of Optix domain and part of Dpp domain. | Some R22D12-Gal4 cells are smaller and likely off-target (see Pxb+ cell in top right corner of Fig. S1D-D', smaller cells also express Dac); these were not counted in lineage trace data. Crosses to lineage tracing lines also reduced intensity of reporter and so smaller numbers of cells were labeled/counted. | Posterior 1/3 of ventral Optix domain, more anterior than Dm4/12 (because Dm1 is not in the Dpp dataset, but Dm4 might be and Dm12 is in the dataset)  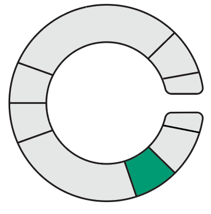 |
| Dm2 (800) | N/A (No LexA line) | Pxb+, Optix+, low Dpp+, Hh+, vOptix+, low dOptix+ (see Notes) | - | scRNAseq data showed robust numbers of Dm2s in Pxb, Optix, Hh and vOptix datasets, but little to no Dpp/dOptix. However, in both dOptix and Dpp lines (Fig. S2), a large number of Im12 cells form a trajectory connected to the Dm2 cells: many Dm2 cells were probably annotated as immature neurons cluster 12. | Entire OPC  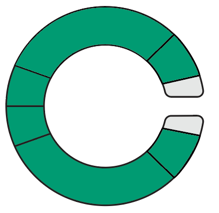 |
| Dm3a (cell number unknown) | Pxb+, Optix+, Hh+ (labels all Dm3 neurons) | Optix+, Hh+, vOptix+  FigS2: Dm3a forms a trajectory with Im11 in the Pxb library, Dm3b forms a trajectory with Im11 in the Dpp libraries and the dOptix 1 library.  It is therefore likely that Dm3a/b cells are mostly annotated as Im11 in the Pxb, Dpp and dOptix datasets. | - | *Act>>lacZ* lineage trace unable to distinguish between Dm3a/b, some R20D11 labeling likely to be off target (e.g. some “Dm3” cells inappropriately express Dll).  The production of Dm3a/b in Pxb, dOptix and Dpp domains is not certain.  Because Dm3a is more clearly found in the Pxb dataset and Dm3b in the dpp dataset, we hypothesize that Dm3a is produced more anteriorly and Dm3b more posteriorly. | vOptix, possibly dOptix/Pxb/Dpp  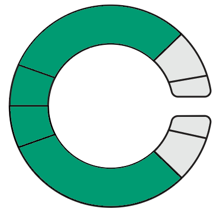 |
| Dm3b (cell number unknown) | Pxb+, Optix+, Hh+ (labels all Dm3 neurons) | Optix+, Hh+, vOptix+  FigS2: Dm3a forms a trajectory with Im11 in the Pxb library, Dm3b forms a trajectory with Im11 in the Dpp libraries and the dOptix 1 library.  It is therefore likely that Dm3a/b cells are mostly annotated as Im11 in the Pxb, Dpp and dOptix datasets. | - | *Act>>LacZ* lineage trace unable to distinguish between Dm3a/b, some R20D11 labeling likely to be off target (e.g. some cells inappropriately express Dll).  The production of Dm3a/b in Pxb, dOptix and Dpp domains is not certain.  Because Dm3a is more clearly found in the Pxb dataset and Dm3b in the Dpp dataset, we hypothesize that Dm3a is produced more anteriorly and Dm3b more posteriorly. | d/v Optix, Dpp  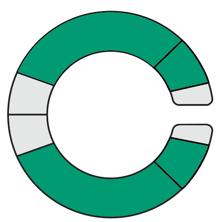 |
| Dm4 (40) | Pxb-, Optix+, Hh+ | Low Optix+, some Dpp+ (1 of 2 libraries), Hh+, vOptix+ | SoxN+Tj+ (which corresponds to Dm1, 4 and 12) = posterior 1/3 of Optix domain and Dpp domain. | All data shows Dm4s born from ventral Optix region. No immature Dm4s found in Dpp datasets, but some Dm4 might have been misidentified as Dm12 (they are very similar as shown by their proximity on UMAP) | Posterior 1/3 of ventral Optix domain, likely at border of Dpp/Optix expression (because Dm1 is not in the Dpp dataset, but Dm4 might be and Dm12 is in the dataset.)  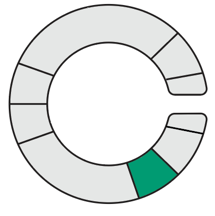 |
| Dm6 (35) | Pxb-, Optix-, Hh- | N/A (no cluster) | Markers: Ap-, Bsh-, Runt-, Lim3-, Dfr-, Hbn-, Dac-, Otd-, Toy-, Dll-, **Svp+**, Scro-, Ey-, vGlut-, Tj-. | Markers suggest that Dm6/17 are possibly contained within cluster 21, which is mostly Wg+ (with very low numbers of cells represented in Dpp, vOptix, Hh, libraries; Rana el-Danaf et al., in preparation). R38H06-Gal4 is very bright and can have bleed-through in neighboring channels, so we disregarded any lines that labeled nuLacZ along with the entire cell body. | Dorsal Wg  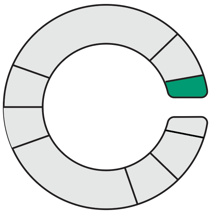 |
| Dm8 (800) | Pxb-, Optix+, Hh+ | Some Pxb+, Optix+, Hh+, vOptix+ | Dac+Tj+ cells in anterior 2/3 of Optix domain and posterior 1/3 of dorsal Optix domain, very small numbers in Pxb domain as well. | Some Dm8s detected in dOptix dataset at the base of the Dm11 and Dm9 trajectories (likely to be misannotated Dm9 or Dm11, or DRA-Dm8: Courgeon and Desplan 2019 -lineage trace shows dorsal Optix origin). Remaining cells are Optix and vOptix/Hh. Small numbers of Pxb cells in Dm8 scRNAseq dataset, and small numbers of Dac+Tj+ cells sit in Pxb region so it is possible that some pDm8s are born from Pxb. | Anterior 2/3 of ventral Optix domain (p/yDm8), posterior 1/3 of dorsal Optix domain not within Dpp region (DRA-Dm8)  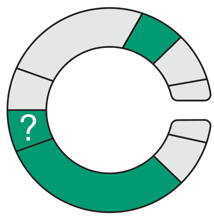 |
| Dm9 (110) | Very few Pxb+, Optix+, very few Hh+ (some Dm9 also labeled as Hh with different LexAOp>>myrRFP lineage trace) | Low Optix+, many Dpp+, Hh+, vOptix+, dOptix+ | Otd+Dve+ Dm9 cells mostly cluster at anterior 2/3 of dorsal Optix region, with a very small cluster in the anterior 1/3 of the ventral Optix domain (see Fig. S5B’). A small population of cells is also visible in Pxb. In more lateral slices, there is no overlap with Dpp; in more medial slices, Dpp clearly overlaps with Dm9 (see Fig. S5A-A’). | Likely that Dm9 scRNAseq cluster is mix of multiple types. Dve is a specific marker of Dm9 at P15 according to scRNAseq, yet multiple groups of cells are labeled *in vivo*. Most cells reside within the anterior 2/3 of dorsal Optix, so that is its likely origin. | Anterior 2/3 of ventral Optix domain (+ posterior Optix/Dpp domain in medial optic lobe).  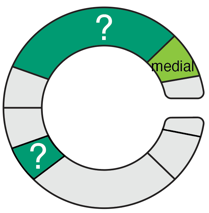 |
| Dm10 (300) | N/A (No LexA line) | Optix+, low Dpp+, Hh+, vOptix+ | Tup+Toy+Ey+ cells cluster in ventral Optix+Dpp region across the whole domain | Between immunostain and scRNAseq, Tup+Toy+Ey+ cells sit across entire vOptix/Dpp regions. | Ventral Optix + Dpp regions  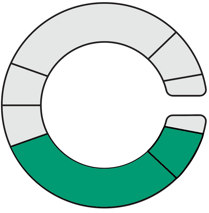 |
| Dm11 (90) | Very few Pxb+, Optix+, Hh- | Pxb+ (but low confidence in the annotation), Optix+, Dpp+, dOptix+ | Dac+Tj+ dorsal cells label posterior 1/3 of dorsal Optix domain + some cells in dorsal Dpp domain | Pxb+ cells likely label ventral Optix Dac+Tj+ cells of unknown identity (see Dm8).  In the Pxb dataset, a trajectory is formed between GMC3 (immature), Dm11 and Dm8, and Dm11 are annotated with low confidence (Fig S2). It is therefore possible that Pxb+ Dm11 cells are in fact misannotated Dm8 cells. | Posterior 1/3 of dorsal Optix region overlapping with Dpp  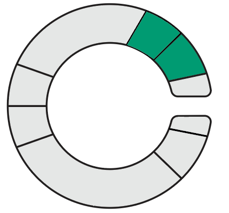 |
| Dm12 (120) | Very few Pxb+, Optix+, Hh+ | Optix+, Dpp+, Hh+, vOptix+ | SoxN+Tj+ (which corresponds to Dm1, 4 and 12) = posterior 1/3 of Optix domain and Dpp domain. | Dm12 well-represented in vOptix and Dpp datasets as well as lineage tracing lines. | Posterior 1/3 of ventral Optix domain at region where Dpp+Optix overlap (because Dm1 is not in the Dpp dataset, but Dm4 might be and Dm12s are).  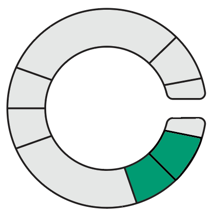 |
| Dm13 (10) | Pxb+, Optix-, Hh+ | N/A (no cluster) | Markers: Ap-, Bsh-, Runt-, **Lim3+**, Dfr-, Hbn-, Dac-, Otd-, Toy-, Dll-, Svp-, Scro-, **Ey+,** Slp-, D-, Tj-, vGlut- | Only a handful of cells labeled with Pxb (2/10), so is possible that these were off-target and that Dm13 is born from Wg. | Ventral Pxb, but possibly ventral Wg.  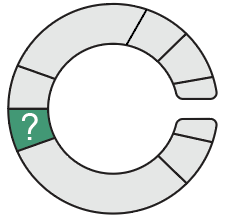 |
| Dm14 (10) | N/A (no LexA line) | N/A (no cluster) | Markers: Ap-, Bsh-, Runt-, **Lim3+**, Dfr-, Hbn-, Dac-, Otd-, Toy-, Dll-, Svp-, Scro-, Ey-, Slp-, D-, Tj-, **vGlut+** | Cluster too small to be reliably identified in dataset. | Unknown |
| Dm15 (250) | Pxb+, Optix+, Hh- | Pxb+, Optix+ | Markers: **scro+, VGlut+,** **Vsx2+,** ap-, svp-, ey-, dac-, oc-, toy-, Dll-, bsh-, run-, Lim3-, vvl-, tj- | Cluster 124 expresses all known Dm15 markers except for DIPδ/DIPθ (Cosmanescu et al., 2018). Possible that Cluster 124 expresses DIPδ/DIPθ at sufficiently low levels that they are hard to detect in dataset. In figures where cluster 124 data is used, the cluster will be labeled “Dm15?”  Dm15? are clearly Optix+ and vOptix-, since these datsets were obtained in adults. They are therefore likely dOptix+, and Dm15? cells are found in dOptix library 2 and form a trajectory with immature neurons 11 (which are also found in dOptix library 1, FigS2). | Dorsal Pxb + Optix  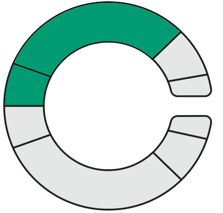 |
| Dm16 (100) | N/A (no LexA line) | N/A (no cluster) | Markers: Ap-, Bsh-, Runt-, Lim3-, Dfr-, Hbn-, Otd-, Toy-, Dll-, Svp-, **Scro+**, Ey-, Tj-, **vGlut+** | No cluster found that expresses same combination of genes. Possibly within a mixed cluster. | Unknown |
| Dm17 (5) | Pxb-, Optix-, Hh- | N/A (no cluster) | Markers:  Ap-, Bsh-, Runt-, Lim3-, Dfr-, Hbn-, Dac-**,** Otd-, Toy-, Dll-, **Svp+**, **Scro+**, Ey-, Tj-, **vGlut+** | Dm17 cell body is ~10μm, so smaller (<2μm) cell bodies were not counted; markers suggest that Dm6/17 are possibly contained within cluster 21, which is mostly Wg+ (with very low numbers of cells represented in Dpp, vOptix, Hh, Rana el-Danaf et al., in preparation) | Dorsal Wg  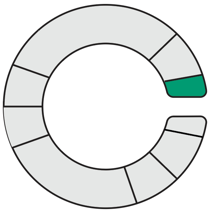 |
